# WNT5A-RHOA axis is a new vulnerability in small-cell lung cancer

**DOI:** 10.1101/2022.04.14.488408

**Authors:** Kee-Beom Kim, Dong-Wook Kim, Youngchul Kim, Jun Tang, Nicole Kirk, Yongyu Gan, Bingliang Fang, Jae-Il Park, Yi Zheng, Kwon-Sik Park

**Affiliations:** Department of Microbiology, Immunology, and Cancer Biology, University of Virginia, Charlottesville, VA 22908, USA; Department of Biostatistics and Bioinformatics, Moffitt Cancer Research Center, Tampa Bay, FL 33612, USA; Department of Thoracic and Cardiovascular Surgery, MD Anderson Cancer Center, Houston, TX 77030, USA; Department of Experimental Radiation Oncology, MD Anderson Cancer Center, Houston, TX 77030, USA; Devision of Experimental Hematology and Cancer Biology, Cincinnati Children’s Hospital Medical Center, Cincinnati, OH 45229, USA

## Abstract

WNT signaling presents an attractive target for cancer therapy due to its widespread oncogenic role. However, the molecular players involved in WNT signaling and the impact of their perturbation remain unknown for numerous recalcitrant cancers including small-cell lung cancer (SCLC). Here we show that beta-catenin, a master mediator of canonical WNT signaling, is not required for SCLC development in genetically engineered mouse models (GEMMs) and its transcriptional program is largely silenced during tumor development. Instead, inactivation of p130 in SCLC cells induces expression of WNT5A, a ligand for beta-catenin-independent WNT pathways. WNT5A is both sufficient and required for SCLC development and cell proliferation and selectively induces *Rhoa* transcription and activates RHOA protein to drive SCLC. *Rhoa* knockout suppresses SCLC development in vivo, and chemical perturbation of RHOA selectively inhibits SCLC cell proliferation. These findings suggest a novel requirement for the WNT5A-RHOA axis in SCLC that is distinct from other noncanonical WNT pathways. This vulnerability of p130-WNT5A-RHOA pathway provides critical insight into the development of novel therapeutic strategies for the recalcitrant cancer, as well as the stratification of patients who may benefit from them. This study also sheds new light on the heterogeneity of WNT signaling and the molecular determinants of its cell-type specificity.

## Introduction

A number of tumors are driven by loss of function mutations in tumor suppressor genes, resulting in few actionable molecular targets for therapeutic intervention. This problem is particularly relevant to small-cell lung cancer (SCLC), an exceptionally lethal disease with a 5-year survival rate of less than 7% that accounts for 13-15% of all lung cancers but results in approximately 30,000 deaths each year in the United States (*1, 2*). This high mortality rate is due to invariable resistance to chemo/radiation therapies and limited efficacy of current immunotherapies, and targeted therapies thus remain an unmet need (*1, 3*). SCLC development is mostly driven by loss-of-function alterations in tumor suppressors, including the near-universal inactivation of RB and P53 and frequent loss of CBP/p300 (*4-7*). While not directly actionable, these alterations cause widespread transcriptional changes, including in developmental signaling pathways such as the WNT pathway, which can be explored as alternative targets (*8, 9*).

Aberrant activity of the beta-catenin-dependent ‘canonical’ WNT pathway drives the development and progression of numerous cancers, including non-small cell lung cancer (NSCLC), breast cancer, and colon cancer. In these cancers, inactivating mutations in regulators such as APC and AXIN2 can increase beta-catenin expression and co-activator action on the TCF/LEF1 family of transcription factors, leading to abnormally high levels of MYC, CCND1, and other proteins that drive proliferation (*10-12*). However, several cancer types, such as sarcoma, do not require WNT/beta-catenin pathway activity for their pathogenesis, suggesting context-dependent functions for this pathway rather than a universally oncogenic role (*8, 9, 13-16*). Further, growing evidence suggests an oncogenic role for beta-catenin-independent ‘non-canonical’ WNT pathways, including WNT/PCP (planar cell polarity) and WNT/Ca^2+^ pathways (*12, 17-19*). In these pathways, small GTPases (RHO/RAC1/CDC42), PI3K, and PLCG1 mediate signals triggered by WNT ligand-receptor interactions and activate downstream effectors such as ROCK1, AKT, and MAP Kinases (JNK, p38, ERK1/2). These effectors in turn regulate actin cytoskeleton homeostasis, cell adhesion, and other functions influencing cell migration, polarity, proliferation, and differentiation (*20-22*).

Both beta-catenin-dependent and -independent WNT pathways play important roles not only in lung development and regeneration but also in lung malignancies (*23-26*). Deregulation of the WNT/beta-catenin pathway promotes lung adenocarcinoma (*24, 27, 28*), while activation of the beta-catenin-independent pathway by WNT5A promotes proliferation of human bronchial epithelial cells transformed by cigarette smoke, as well as cell invasion and proliferation of lung cancer cells (*29, 30*). Additional studies have found that NSCLC dependence on WNT pathway activity shifts between the canonical and non-canonical pathways over the course of lung cancer progression (*31*). Less is known of the role of WNT signaling in SCLC pathogenesis. A microarray analysis has shown down-regulation of the WNT/beta-catenin pathway in SCLC (*32*), while more recently, whole-exome sequencing revealed this pathway to be active in relapsed SCLC tumors following standard chemotherapy (*33*). WNT11, a ligand for the beta-catenin-independent pathway, has been shown to promote proliferation of SCLC cells (*34*). However, these studies do not consistently link the two WNT pathways nor do they conclusively determine whether either plays an oncogenic or tumor-suppressive role.

In this study we aim to characterize the WNT pathway in SCLC and determine its physiological role using genetically engineered mouse models (GEMMs). In these models, adenovirus Cre (Ad-Cre)-mediated deletion of *Rb* and *p53* in the lung epithelium induces lung tumors that closely recapitulate the major histopathological features of human SCLC (*35*). This study also involves genetic and chemical perturbations in primary tumor cells and precancerous neuroendocrine cells (preSCs) derived from the GEMMs (*36*). PreSCs in particular allow for efficiently testing of candidate drivers for the ability to transform precursor cells to SCLC (*36-39*). Our data demonstrate that beta-catenin is not active in, and dispensable for, SCLC development. Instead, activation of the beta-catenin-independent WNT pathway by WNT5A promotes tumorigenic transformation of preSCs and is required for SCLC development and cell proliferation. We also demonstrate that loss of p130 induces *Wnt5a* expression, and WNT5A induces RHOA expression and depends on RHOA activity to drive cell proliferation. These findings suggest the existence of a WNT pathway unique to SCLC and present actionable targets for tumor intervention.

## Results

### Beta-catenin is not required for SCLC development and cell proliferation

To begin mapping out WNT pathway activity in SCLC development *in vivo*, we crossed *Rb*/*p53*-mutant (*RP*) mice with *Axin2*^+/lacZ^ mice. This *Axin2*-*lacZ* strain expresses the reporter gene encoding beta-galactosidase under the control of the *Axin2* promoter (*40*). Given that X-gal staining for visualizing beta-galactosidase expression parallels AXIN2 expression, and *Axin2* is a canonical transcriptional target of the WNT/beta-catenin pathway, this system generally reports WNT pathway activity in various tissues, including intestinal crypts (Supplementary Fig. 1) (*40*). Nine months following infection with Ad-Cre, the lungs of *RP Axin2*^+/lacZ^ mice showed strong X-gal staining in nodular tumors and small lesions, but not in non-tumor areas (Fig. 1A). X-gal staining in lung tumors overlapped with immunostaining for SYP, a protein marker of SCLC (Fig. 1B, Supplementary Fig. 2A). X-gal staining was also detected in metastatic tumors in the liver of *RP Axin2*^+/lacZ^ mice (Supplementary Fig. 2B), and in primary cells derived from lung tumors (Supplementary Fig. 2C), and subcutaneous tumors generated from these cells (Supplementary Fig. 2D). However, we did not observe X-gal staining in pulmonary neuroendocrine cells, the main cell type of origin for SCLC (Supplementary Fig. 1) (*41*).

**Fig. 1.**
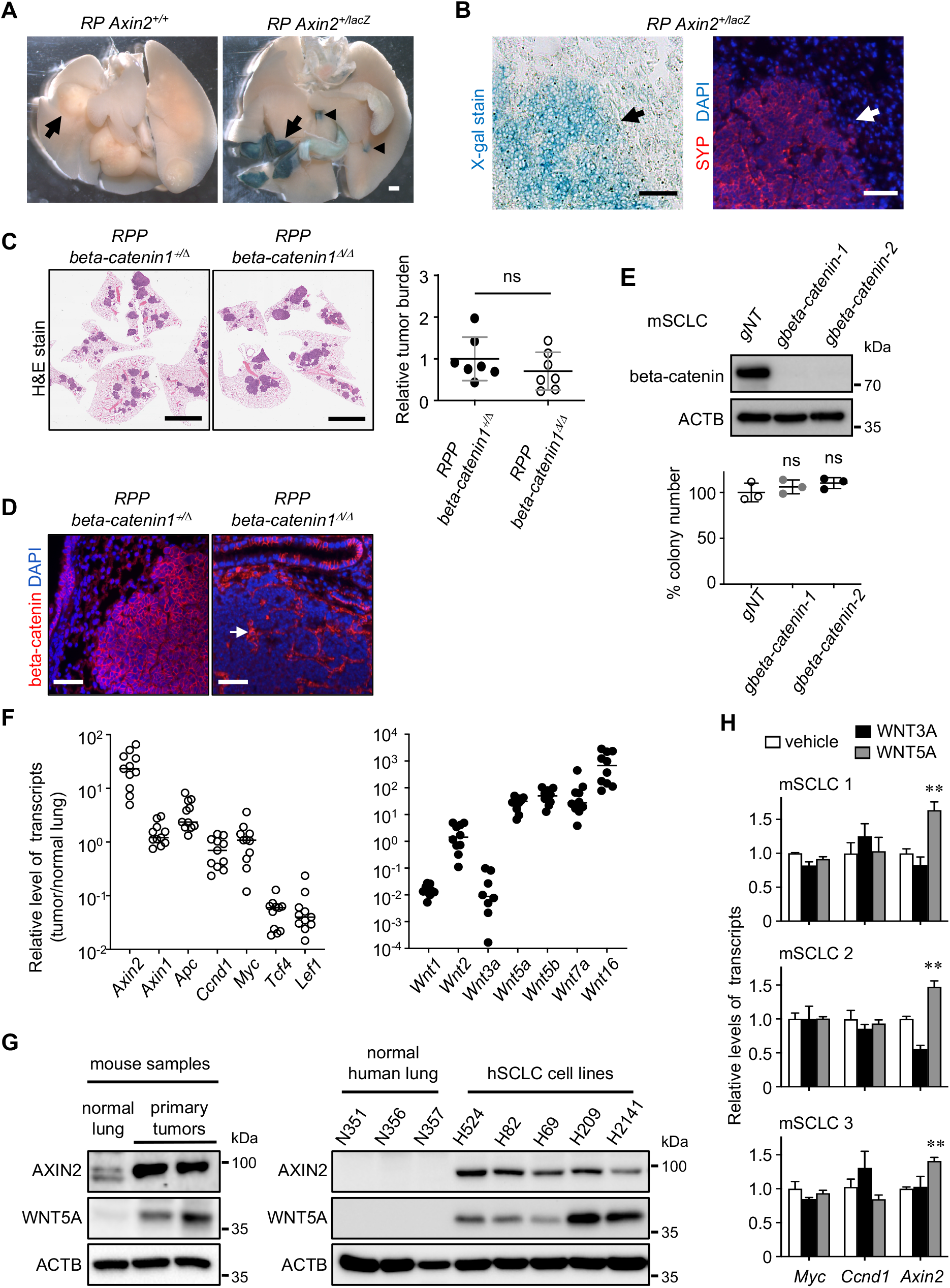
Beta-catenin is not required for SCLC development. **(A)** Whole-mount X-gal stained lungs from *Rb/p53 (RP) Axin2*^*+/+*^ and *Axin2*^*+/lacZ*^ mice 9 months after Ad-Cre infection. The interior of cut lungs is shown, and arrows and arrowheads indicate tumors and small lesions, respectively. **(B)** Representative sections containing X-gal stained lung tumors and adjacent tissue, visualized using light microscopy (left) and immunofluorescent staining for SYP (red) marking neuroendocrine cells (right). DAPI (blue) was used as a counterstain. Arrows indicate normal tissues either positive or negative for X-gal and SYP staining. **(C)** Representative images of H&E-stained lung sections from *Rb/p53/p130*-mutant (*RPP*) mice with additional *beta-catenin*^+/lox^ or *beta-catenin*^lox/lox^ alleles (left) and quantification of tumor burden (tumor area/lung area, n=7 per genotype) (right). **(D)** Immunostaining for beta-catenin and DAPI on lung sections. Arrow indicates beta-catenin expression in stroma cells. **(E)** Immunoblot for beta-catenin in *beta-catenin*-targeted mouse SCLC (mSCLC) cells (top), and quantification of soft agar colonies formed from *beta-catenin-*targeted and control cells (bottom) (n=3 per cell type). **(F)** Quantitative PCR analysis measuring the levels of transcripts encoding the ligands and other components of the WNT/beta-catenin pathway in primary tumors relative to normal lung. **(G)** Immunoblots for WNT5A and AXIN2 in mouse primary tumors and human SCLC lines compared with normal lung tissue. ACTB was used a loading control. **(H)** Quantitative PCR analysis measuring the transcripts of *Myc, Ccnd1*, and *Axin2* in multiple lines of mSCLC cell lines treated with vehicle, WNT3A peptide, or WNT5A peptide (100 ng/mL). *, *p*<0.01; **, *p*<0.001. Statistical tests were performed using unpaired t-test (ns: not significant). Error bar represents standard deviation. Scale bars: A, 1 mm; B, 50 μm; C, 5mm; D, 50 μm.

Since the *Axin2* reporter, and by extension the WNT/beta-catenin pathway appears to be activated specifically in tumor lesions and tumor cells, we next sought to better understand the role of this pathway in SCLC by conditionally deleting *beta-catenin* in *Rb*/*p53*/*p130*-mutant (*RPP*) mice. In this mouse model, Ad-Cre-mediated deletion of floxed alleles of *p130* (an *Rb* homolog) as well as *Rb* and *p53* in lung epithelial cells results in more than a dozen nodular tumors with a latency of 6 months, serving as a robust model for testing the tumor-suppressive effect of genetic perturbations (*36, 42, 43*). Tumor burden (tumor area/lung area) was not significantly different in the lungs of *RPP beta*-*catenin*^+/lox^ vs *RPP beta-catenin*^lox/lox^ mice infected with Ad-Cre (Fig. 1C), although staining confirmed beta-catenin loss (Fig. 1D). Furthermore, we tested whether the WNT/beta-catenin pathway is required for SCLC proliferation, using CRISPR-mediated targeting of *beta-catenin* in mouse SCLC (mSCLC) cells derived from *RP* mice (*42*). These *beta-catenin*-targeted cells formed colonies in soft agar at a rate similar to non-targeted control cells (Fig. 1E). These results suggest that beta-catenin is not required for SCLC development or for the continuing expansion of tumor cells, but are seemingly inconsistent with the presence of active WNT/beta-catenin pathway as indicated by *Axin2* reporter activity. However, while *Axin2* transcript levels were higher in tumors than normal lung tissue, consistent with reporter activity, expression levels of other transcriptional targets or pathway components of the WNT/beta-catenin pathway were lower (*Ccnd1, Myc, Tcf4*, and *Lef1*) or not significantly different (*Axin1* and *Apc*) (Fig. 1F). These results suggest that *Axin2* expression is induced in SCLC tumors in the absence of an active WNT/beta-catenin pathway.

We hypothesized that other regulators of *Axin2* expression may be implicated in SCLC development, and focused on WNT5A, a ligand for beta-catenin-independent WNT pathways that can nevertheless induce *Axin2* expression in certain contexts (*44, 45*). RT-qPCR analysis showed that genes encoding WNT ligands for beta-catenin-independent pathways (*Wnt5a, Wnt5b, Wnt7a*, and *Wnt16*) were upregulated in tumors vs normal lung tissue (Fig. 1F). Conversely, transcripts encoding ligands for the canonical WNT/beta-catenin pathway, including *Wnt1, Wnt2*, and *Wnt3a*, were either not significantly enriched or absent in tumors (Fig. 1F). Immunoblot analysis validated the differential expression patterns of AXIN2 and WNT5A between tumors and normal lung tissue in mice, and between human SCLC cell lines and normal lung cells (Fig. 1G). Furthermore, treatment with recombinant peptides revealed that WNT5A, not WNT3A, increased *Axin2* transcript levels in three independent mSCLC cell lines relative to vehicle-treated cells, while *Myc* and *Ccnd1* transcript levels were unchanged following either treatment (Fig. 1H). Taken together, these data establish WNT5A as a driver of *Axin2* expression and a candidate oncogene in SCLC.

### WNT5A promotes tumorigenic progression of preneoplastic cells

We next used a precancerous neuroendocrine cell (preSCs)-based model of SCLC development to continue interrogating the role of WNT5A in SCLC. These preSCs, derived from early-stage lesions developed in SCLC mouse models, remain preneoplastic in culture but transform into SCLC upon activation of oncogenic drivers (*36-39*). Immunoblot analysis revealed that WNT5A and AXIN2 levels were higher in SCLC cells than in preSCs (Fig. 2A). However, preSCs expressing ectopic WNT5A (*Wnt5a*-preSCs) had increased AXIN2 levels compared to control preSCs transduced with empty viral vector (Fig. 2B), and the difference in AXIN2 expression was similar to that seen between mSCLC cells and preSCs (Fig. 2A). Notably, *Wnt5a*-preSCs gave rise to more colonies in soft agar than control preSCs (Fig. 2B), and formed subcutaneous tumors faster than control preSCs when implanted in the flanks of athymic nude mice (Fig. 2C). H&E staining and immunostaining demonstrated that the subcutaneous tumors generated from both *Wnt5a*-preSCs and control preSCs showed well-known features of SCLC, including high nuclear/cytoplasmic ratio and CGRP expression (Fig. 2D). Staining for phosphorylated-histone H3 (pHH3) revealed significantly more mitotic cells in the tumors generated from *Wnt5a*-preSCs compared to control preSCs (Fig. 2E). Taken together, these results suggest that increased expression of WNT5A promotes the neoplastic transformation of preSCs into SCLC.

**Fig. 2.**
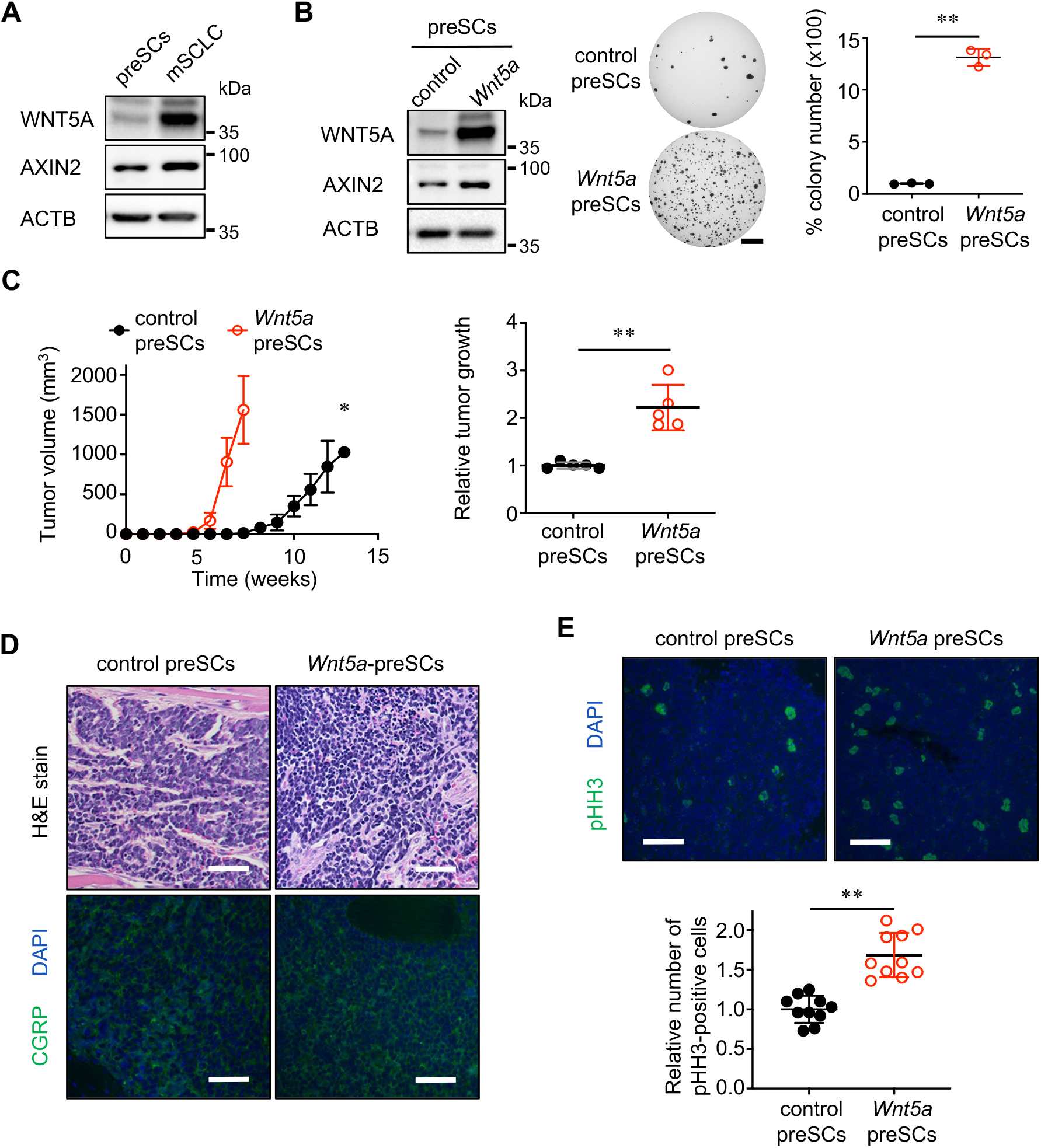
WNT5A induces tumorigenic progression of precancerous cells (preSCs). **(A)** Immunoblot for WNT5A and AXIN2 in preSCs and mSCLC cells. ACTB was used as a protein loading control. **(B)** Immunoblot for WNT5A and AXIN2 in control and *Wnt5a*-preSCs. ACTB was used as a protein loading control (left). Representative images (middle) and quantification of soft agar colonies (>0.2mm) (right) from control and *Wnt5a*-preSCs (n=3 per cell type). **(C)** Quantification of allograft tumors over time, following injection of mice with control and *Wnt5a*-preSC cells (left), and of subcutaneous tumors >1.5 cm in diameter, where relative tumor growth represents tumor weight (g, grams) divided by latency (days after allograft) (right). **(D)** Representative H&E-stained sections of subcutaneous tumors derived from *Wnt5a*-preSCs (top), and immunostaining for CGRP marking neuroendocrine cells (green; bottom). **(E)** Representative images of immunostained lung sections (top) and quantification of pHH3-positive cells per lung area (bottom) in subcutaneous tumors formed from control and *Wnt5a*-preSCs (n=10 tumors per genotype). DAPI (blue) was used as a counterstain. *, *p*<0.01; **, *p*<0.001. Statistical tests were performed using unpaired t-test. Error bar represents standard deviation. Scale bars: B, 5 mm; D, E, 50 μm.

### Inactivation of p130 induces expression of WNT5A in SCLC

Given the importance of WNT5A in neoplastic transformation, we sought to better understand the mechanisms underlying WNT5A regulation. The transcriptional repressor, E2F4, is known to occupy the regulatory element of *Wnt5a* in neural precursor cells in conjunction with p130 and the other Rb family member p107 *(46, 47*). We therefore postulated that the repressor function of E2F4 is weakened upon p130 inactivation, which is frequent in SCLC (*5*), and its targets are transcriptionally de-repressed (Fig. 3A). To test this idea, we examined the impact of knocking out *p130* in preSCs using CRISPR/Cas9. *p130*-*ko* preSCs showed a drastic reduction in p130 levels and induction of WNT5A expression (Fig. 3B), coinciding with an increase in *Wnt5a* transcript levels (Fig. 3B). This increased WNT5A expression was maintained in subcutaneous tumors derived from *p130*-*ko* preSCs (Fig. 3C). Consistently, lentiviral restoration of p130 expression in *p130*-*ko* preSCs markedly reduced WNT5A levels (Fig. 3D). To determine if this relationship between p130 and WNT5A exists in spontaneous SCLC tumors, we surveyed primary cells derived from tumors developed in *RP, RPP*, and *RPM* mice that express heterogeneous levels of p130. Similar to *RP* and *RPP* mice, *RPM* (*Rb*/*p53*/*Myc*-mutant) mice develop lung tumors after Cre-mediated induction of a hyperactive mutant allele of *Myc* as well as loss of *Rb* and *p53*, and the tumor cells recapitulate the features of a subset of human SCLC lines used above (*48*). WNT5A protein level was drastically higher in primary cells derived from *RPP* and *RPM* tumors lacking p130 and 107, compared to that in preSCs and primary cells derived from *RP* tumors expressing these proteins (Fig. 3E). Notably, a high level of WNT5A was detected in the *RP* cells with p130 phosphorylation (Fig. 3E), which inhibits the tumor suppressor function (*46*). These findings indicate that WNT5A is induced in SCLC due to the loss or inactivation of p130 and p107, which otherwise repress *Wnt5a* transcription.

**Fig. 3.**
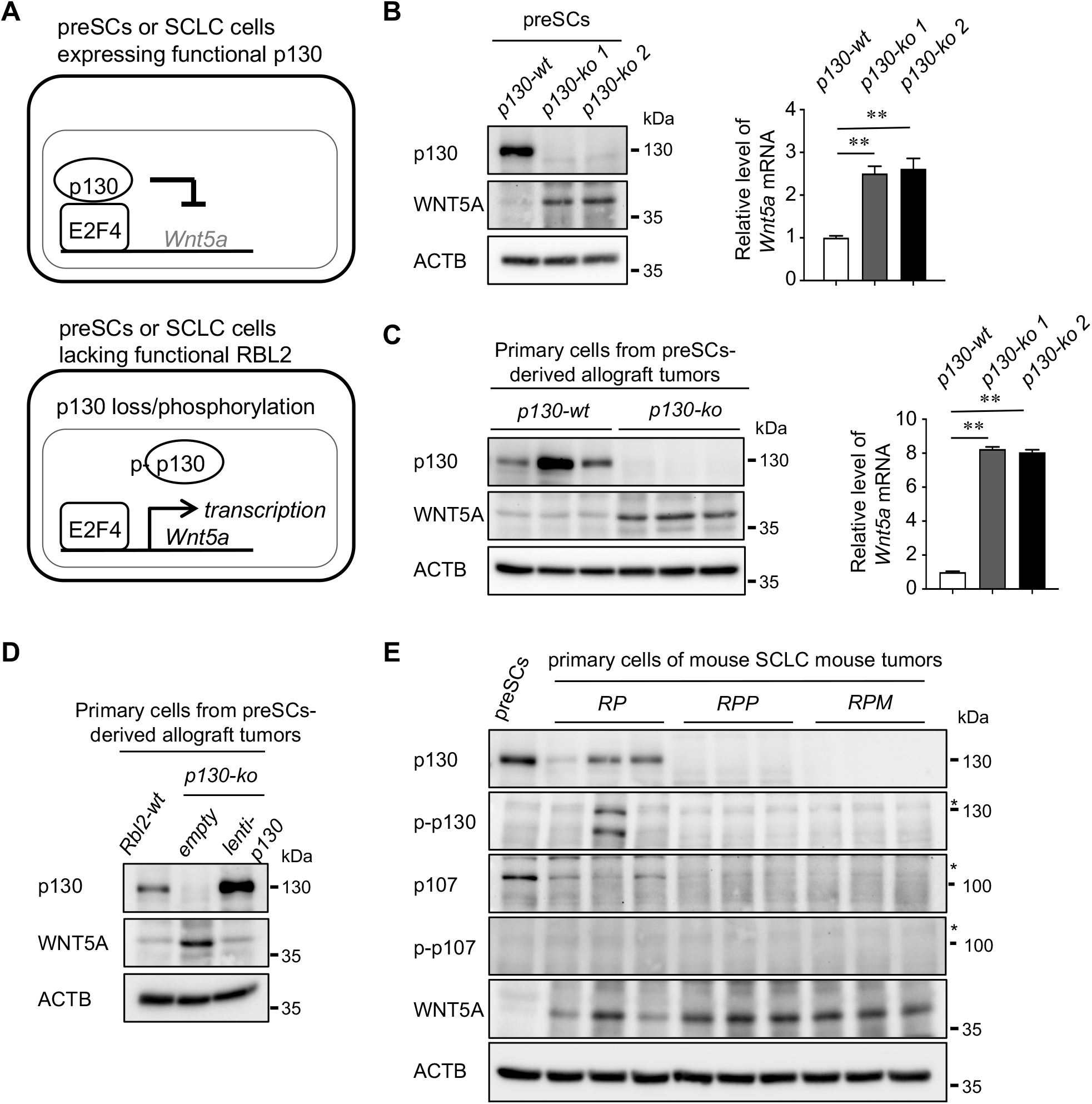
p130 loss induces WNT5A expression. **(A**) A hypothetical model in which E2F4-driven *Wnt5a* transcription is suppressed by the p130-E2F4 repressor complex; in the event that p130 is deactivated by phosphorylation, or lost, the repressor function of E2F4 is weakened and its targets are transcriptionally de-repressed. **(B, C)** Immunoblot for WNT5A and p130 in control and *p130-ko* preSCs and primary cells isolated from preSC-derived allograft tumors (left). RT-qPCR for *Wnt5a* transcript levels normalized to *Gapdh* (n=3 replicates per cell type) (right). **(D)** Immunoblot for WNT5A and p130 in the cells of allograft tumors derived from *p130-wt* and *p130-ko* preSCs, as well as *p130-ko* preSCs infected with lentivirus to rescue p130 expression (*lenti-p130*). **(E)** Immunoblot for WNT5A, p130, p107, and phosphorylated forms of p130 and p107 (p-p130 and p-p107) in primary tumors developed in *RP, RPP, RPM* mice. Asterisk (*) indicates bands with correct sizes. ACTB was used as a protein loading control. Statistical tests were performed using unpaired t-test. Error bar represents standard deviation **, *p*<0.001.

### WNT5A is required for SCLC development and cell proliferation

To determine the necessity of WNT5A for tumor development *in vivo*, we deleted *Wnt5a* in *RPP* mice carrying floxed alleles of the gene, and 6 months after intratracheal Ad-Cre instillation, observed significantly lower tumor burden and fewer mitotic cells in the lungs of *Wnt5a*^Δ/Δ^ vs *Wnt5a*^+/+^ *RPP* mice (Fig. 4A, B). To determine whether these phenotypes are owing to cell-intrinsic differences, we performed cellular and molecular analyses on primary cells derived from these lung tumors. *Wnt5a*^Δ/Δ^ tumor cells gave rise to fewer colonies in soft agar than *Wnt5a*^+/+^ cells (Fig. 4C), and showed a marked increase in levels of cleaved PARP1 and CASP3 indicating higher rates of apoptosis (Fig. 4D). Immunoblot analysis verified WNT5A loss and revealed a drastic reduction in AXIN2 levels in *Wnt5a*^Δ/Δ^ compared with *Wnt5a*^+/+^ cells, but revealed no significant differences in the levels of beta-catenin, MYC, or transcription factors regulating neuroendocrine differentiation (ASCL1, NEUROD1) (Fig. 4E). The tumors developed in both groups of mice displayed histology and expression of neuroendocrine markers such as CGRP typical of SCLC (Supplementary Fig. 3). These findings indicate that WNT5A loss suppresses SCLC tumor development in part through reduced proliferation and increased cell death.

**Fig. 4.**
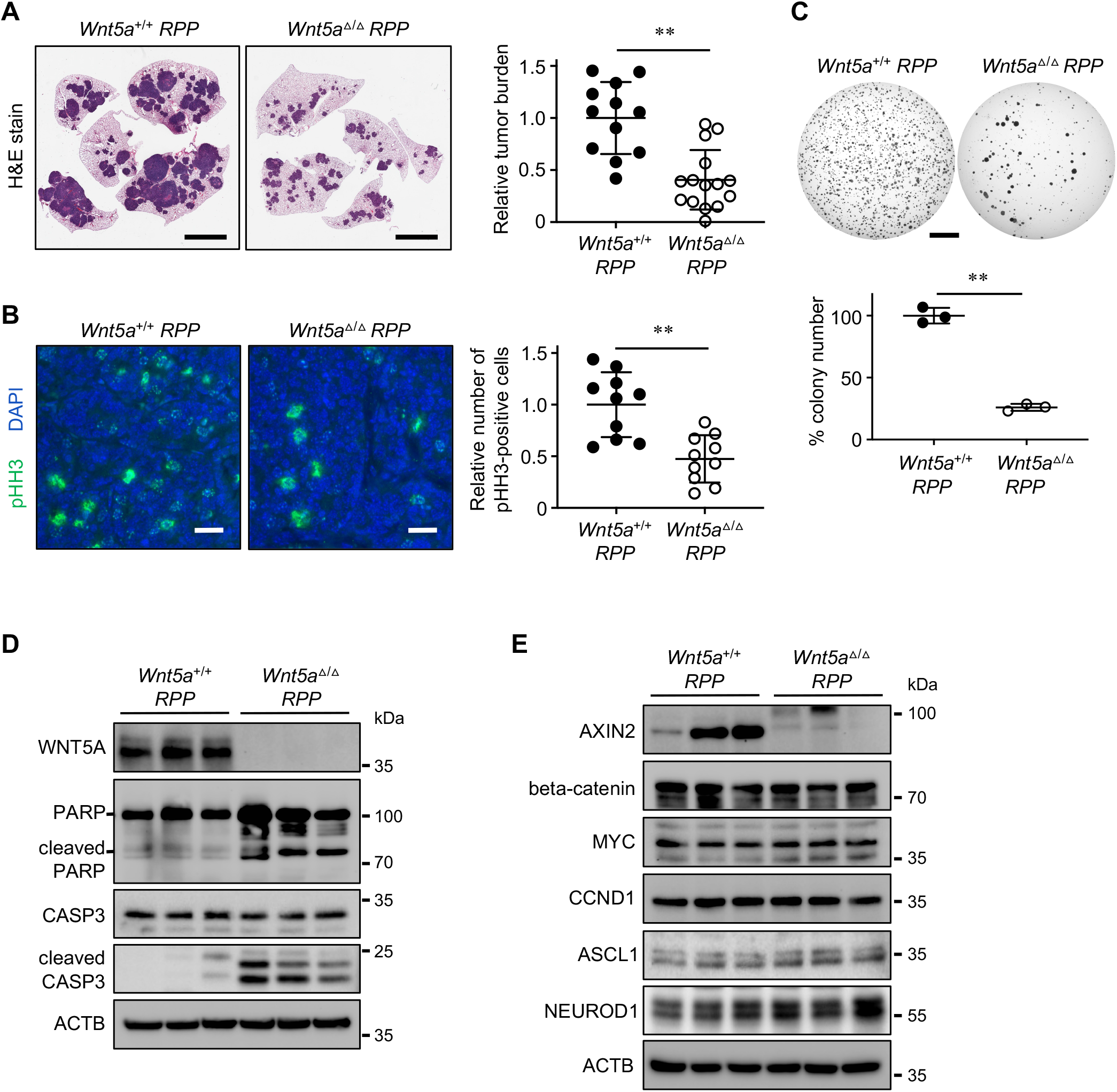
WNT5A loss suppresses tumor development in autochthonous mouse models. **(A)** Representative images of H&E-stained lung sections (left) and quantification of tumor burden (tumor area/lung area) (right) in tumors from *Wnt5a*^*+/+*^ vs. *Wnt5a*^*Δ/Δ*^ *RPP* mice (n=12 and n=15, respectively). **(B)** Representative images of immunostained lung sections (left) and quantification of pHH3-positive cells per lung area (right) in *Wnt5a*^*+/+*^ vs. *Wnt5a*^*Δ/Δ*^ *RPP* mice (n=10 tumors per genotype). DAPI was used as a counterstain. **(C)** Representative images (top) and quantification of soft agar colonies (bottom) formed by primary cells derived from *Wnt5a*^*+/+*^ vs. *Wnt5a*^*Δ/Δ*^ *RPP* tumors (n=3 per cell type). **(D, E)** Immunoblot for cleaved/total CASP3 and PARP1 (D) and beta-catenin-dependent pathway components in *Wnt5a*^*+/+*^ vs. *Wnt5a*^*Δ/Δ*^ *RPP* primary cells. ACTB was used as a protein loading control. **, *p*<0.001. Statistical tests were performed using unpaired t-test. Error bar represents standard deviation. Scale bars: A, C, 5 mm; B, 50 μm.

We next determined the role of WNT5A in the continuing growth of tumor cells by targeting *Wnt5a* in mSCLC cells using CRISPR-mediated gene editing. *Wnt5a*-targeted mSCLC cells gave rise to significantly fewer soft agar colonies than non-targeted control cells (Fig. 5A and Supplementary Fig. 4A), and formed subcutaneous tumors in nude mice at significantly slower rates and with lower proliferation rates indicated by pHH3 positivity (Fig. 5B and Supplementary Fig. 4B). Knocking down *Wnt5a* in human SCLC lines (H69, H82, H524) using lentiviral shRNAs similarly led to reduced colony-forming capacity in soft agar (Fig. 5C, E and Supplementary Fig. 4C), and slower formation of subcutaneous tumors (Fig. 5D), compared with control cells expressing random non-targeting shRNA. Taken together, these results suggest that WNT5A is critical for both the *in vivo* development of SCLC and the continuing expansion of tumor cells.

**Fig. 5.**
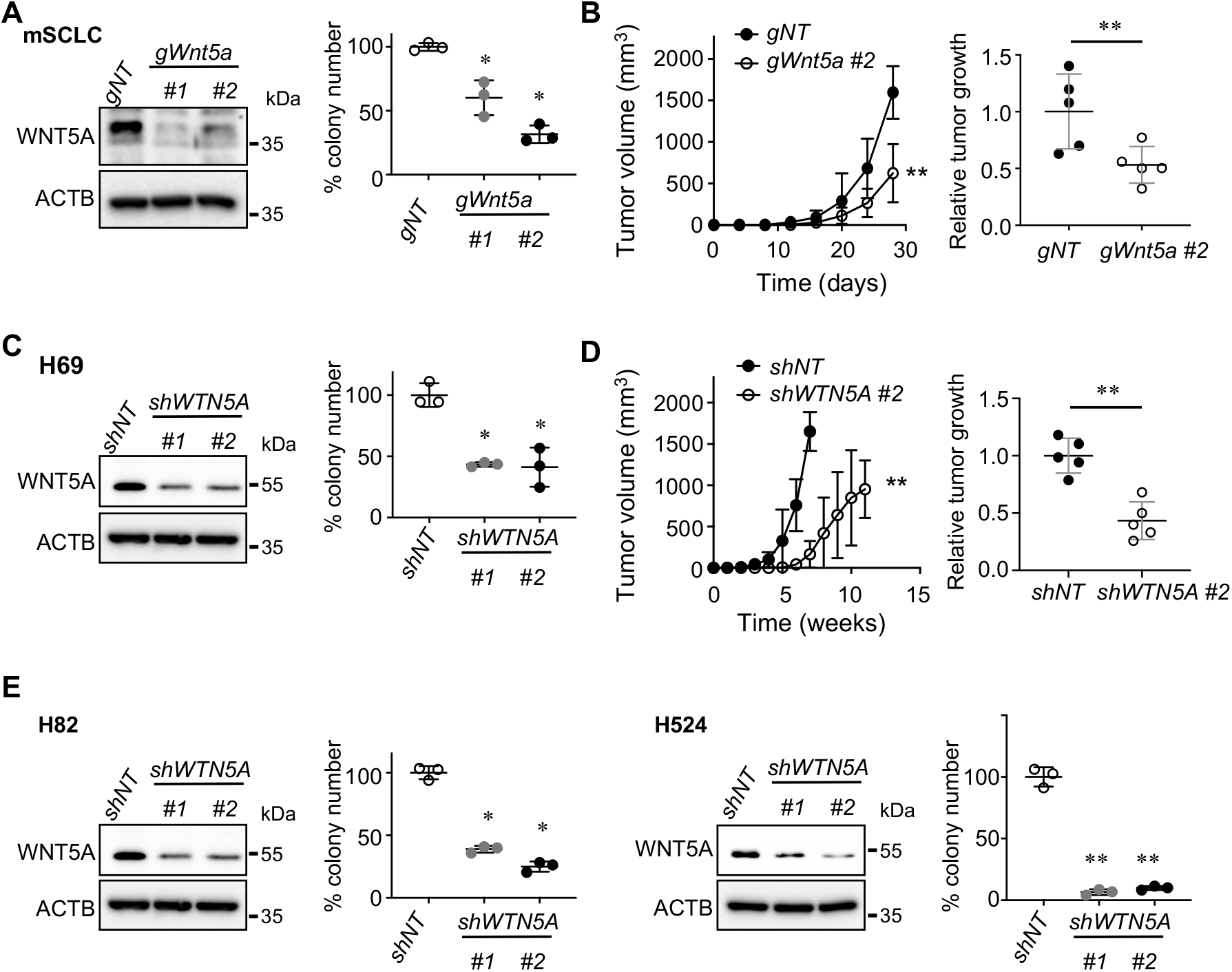
WNT5A is required for the expansion of human SCLC lines and mouse tumor cells. **(A, C, E)** Immunoblot for WNT5A in mSCLC cells derived from tumors developed in *RP* mice, and in human SCLC cell lines (H69, H82, H524) (left). Quantification of soft agar colonies (>0.2 mm in diameter) derived from *Wnt5a*-targeted mSCLC cells, *WNT5A*-knockdown human SCLC cells, and their respective controls (n=3 replicates per cell type) (right). **(B, D)** Volumes of subcutaneous tumors (n=5) generated from the *Wnt5a-*targeted mSCLC cells in (A) and *WNT5A*-knockdown human SCLC cells in (C), along with their respective controls (left). Quantification of subcutaneous tumors >1.5 cm in diameter, where relative tumor growth represents tumor weight (g, grams) divided by latency (days after cell implantation). ACTB was used as a protein loading control. *, *p*<0.01; **, *p*<0.001. Statistical tests were performed using unpaired t-test. Error bar represents standard deviation.

### WNT5A selectively increases expression and activity of RHOA in SCLC

To further characterize the molecular mechanism of WNT5A-driven SCLC tumorigenesis, we took an unbiased approach to identify associated molecular changes using RNA-sequencing. Analysis of RNA-sequencing results showed 821 differentially expressed (DE) genes in *Wnt5a*-preSCs relative to control preSCs (FDR adjusted *p*<0.05), including 495 upregulated and 326 downregulated genes (Fig. 6A, Supplementary Table 1). RT-qPCR analysis validated the increased expression of *Wnt5a, Axin2*, and a handful of other DE genes, while confirming a lack of significant differences in *Myc* and *Ccnd1* expression (Supplementary Fig. 5). Gene set enrichment analysis (GSEA) revealed an enrichment for MYC targets and unfolded protein response (Fig. 6B, Supplementary Table 2).

**Fig. 6.**
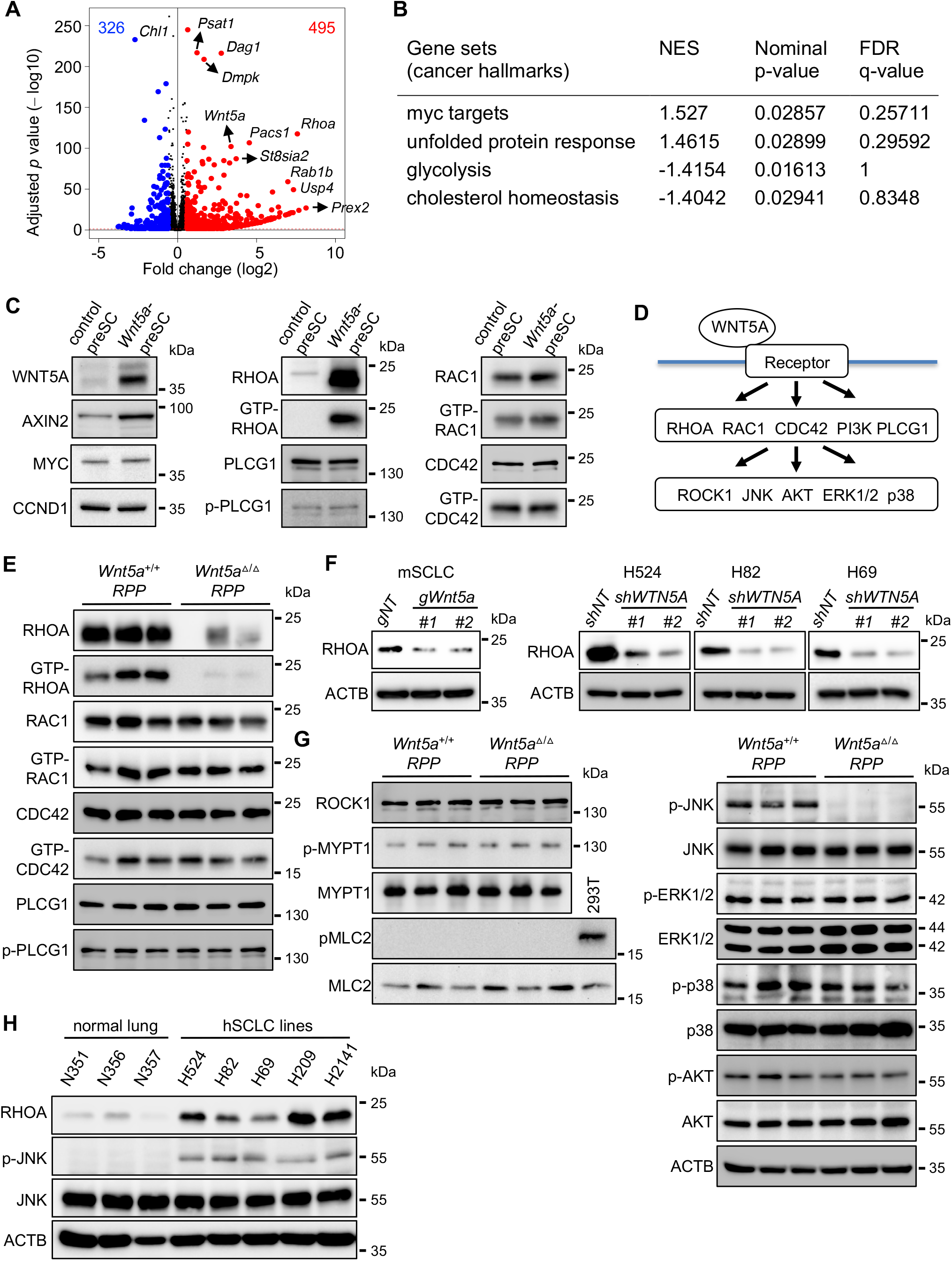
WNT5A selectively induces the expression and activity of RHOA. **(A**) Volcano plot showing differential gene expression between control and *Wnt5a*-preSCs. Red and blue dots (495 and 326, respectively) indicate differentially expressed genes (DEs) with a ≥1.5-fold increase or decrease in expression in *Wnt5a*-preSCs relative to control preSCs; dots above the red horizontal line indicate DEs with significant expression changes at an FDR-adjusted *p* value < 0.05. FDR: false discovery rate. **(B)** Results of gene set enrichment analysis (GSEA), including the gene sets with nominal *p* value <0.05. NES: normalized enrichment score. **(C)** Immunoblot for WNT pathway components in control vs *Wnt5a*-preSCs, including MYC and CCND1 from the CTNNB1-dependent pathway, total and GTP-bound RHOA, PCLG1 from the WNT/Ca^2+^ pathway, and total and GTP-bound RAC1 and CDC42 from the WNT/PCP pathway. **(D)** Diagram depicting potential mediators and effectors of WNT5A signaling pathways. **(E)** Immunoblot for total and GTP-bound RHOA, RAC1, and CDC52, as well as total and phosphorylated PLCG1, in *Wnt5a*^*+/+*^ vs. *Wnt5a*^*Δ/Δ*^ *RPP* cells. **(F)** Immunoblot for RHOA in *WNT5A*-knockdown human SCLC lines. **(G)** Immunoblot for ROCK1 and its substrates (MYPT1 and MLC2) and their phosphorylation status in *Wnt5a*^*+/+*^ vs. *Wnt5a*^*Δ/Δ*^ *RPP* cells (left), and immunoblot for total and phosphorylated (p-) JNK, ERK1/2, P38, and AKT in *Wnt5a*^*+/+*^ vs. *Wnt5a*^*Δ/Δ*^ *RPP* cells (right). **(H)** Immunoblot for RHOA, JNK, and phosphorylated JNK (p-JNK) in human normal lung and SCLC cell lines. ACTB was used as a loading control. *, *p*<0.01; **, *p*<0.001. Statistical tests were performed using unpaired t-test (ns: not significant). Error bar represents standard deviation.

Among the DE genes, *Rhoa* was particularly notable, as it encodes a member of the Rho family of small GTPases known to mediate WNT5A signaling. Consistently, immunoblots showed that WNT5A, AXIN2, and RHOA levels were higher in *Wnt5a*-preSCs than control preSCs, while MYC and CCND1 levels were similar between the two cell types (Fig. 6C). We next examined whether other Rho family members known to mediate WNT5A signaling, including RAC1 and CDC42 in the WNT/PCP pathway and PLCG1 in the WNT/Ca^2+^ pathway, are impacted by WNT5A activity in SCLC (Fig. 6D). We did not observe any significant differences in the levels of RAC1, CDC42 and PLCG1 between control preSCs and *Wnt5a*-preSCs (Fig. 6C), suggesting that RHOA mediation of WNT5A signaling is specific to SCLC. To further examine the relationship between WNT5A and RHOA, we tested the impact of WNT5A loss on RHOA expression in primary cells derived from our mouse models shown in Fig. 3C-E. Levels of RHOA and its GTP-bound ‘active’ form were significantly lower in cells derived from *Wnt5a*^Δ/Δ^ vs *Wnt5a*^+/+^ *RPP* tumors, whereas levels of RAC1 and CDC42 and their GTP-bound forms, and of PLCG1 and its phosphorylated form, were similar between the two cell types (Fig. 6E). Additionally, CRISPR/Cas9-mediated knockout of *Wnt5a* in mouse SCLC cells and shRNA-mediated knockdown of *WNT5A* in human SCLC lines led to a significant reduction in RHOA levels relative to non-targeted cells (Fig. 6F).

We further defined the WNT5A pathway in SCLC by identifying RHOA-associated downstream effectors in mouse SCLC primary cells using immunoblot analysis. ROCK1 activity, measured by phosphorylation levels of a well-known target protein MYPT1, was similar in *Wnt5a*^Δ/Δ^ and *Wnt5a*^+/+^ *RPP* cells (Fig. 6G). Notably, MLC2, another downstream target of the RHOA-ROCK1 axis, was present but not phosphorylated in tumor cells, in contrast to 293T cells (Fig. 6G). We also observed little or no difference in levels of phosphorylated AKT, p38, and ERK1/2, but a marked decrease in phosphorylated JNK in *Wnt5a*^Δ/Δ^ relative to *Wnt5a*^+/+^ *RPP* cells (Fig. 6G). Consistently, RHOA and phosphorylated JNK levels were higher in mouse primary tumors and human SCLC cell lines than their respective normal lung tissue (Fig. 6H). These results suggest that WNT5A signaling in SCLC involves RHOA and JNK and is distinct from the RHO-ROCK axis in the WNT/PCP and WNT/Ca^2+^ pathways.

### RHOA mediates WNT5A signaling pathway in SCLC

To determine the significance of RHOA in WNT5A signaling, we tested if restoration of RHOA using a doxycycline-inducible lentiviral vector would rescue the molecular and cellular phenotypes observed in WNT5A-deficient cells. Expression of a dominant active form of RHOA (RHOA Q63L) in *Wnt5a*^Δ/Δ^ *RPP* tumor cells fully restored its colony-forming ability in soft agar to that of *Wnt5a*^+/+^ *RPP* cells (Fig. 7A and Supplementary Fig. 6A). Next, we measured the effect on SCLC proliferation of blocking RHOA function using Rhosin, a selective inhibitor of GTPase activity among the RHOA subfamily (*49, 50*). Rhosin treatment reduced JNK phosphorylation levels but did not affect the phosphorylation of AKT, p38, and ERK1/2 (Fig. 7B). Rhosin treatment also inhibited the short-term viability of mouse SCLC cells in a concentration-dependent manner, whereas the chemical inhibitors of RAC1 and CDC42 had no effect at a range of concentrations up to 100 μM (Fig. 7C). The treatment also reduced the number of soft agar colonies formed from mouse SCLC cells and *Wnt5a*-preSCs (Fig. 7D and Supplementary Fig. 6B, C). Notably, the inhibitory effect on colony-forming ability was larger for *Wnt5a*-preSCs, which show increased RHOA expression, than control preSCs (Fig. 7D and Supplementary Fig. 6C). Similar effects on viability and soft agar colony formation were seen in human SCLC lines representing multiple molecular subtypes of SCLC (51), except for H526 (Fig. 7E, F and Supplementary Fig. 6D). However, Rhosin did not affect the growth of NSCLC lines at the same doses tested in SCLC cells (Fig. 7E, F). To determine the factors underlying this difference, we examined RHOA expression and activity in SCLC and NSCLC cells using immunoblot. While the levels of total and GTP-bound RHOA varied across cell lines, the ratio of GTP-bound to total RHOA was higher in the Rhosin-sensitive SCLC cells than in H526 and the NSCLC cell lines that did not respond to the drug (Fig. 7G). Notably, WNT5A expression correlated with GTP-RHOA levels and degree of Rhosin sensitivity (Fig. 7G). Taken together, these data suggest that RHOA is induced and activated by WNT5A and promotes SCLC proliferation.

**Fig. 7.**
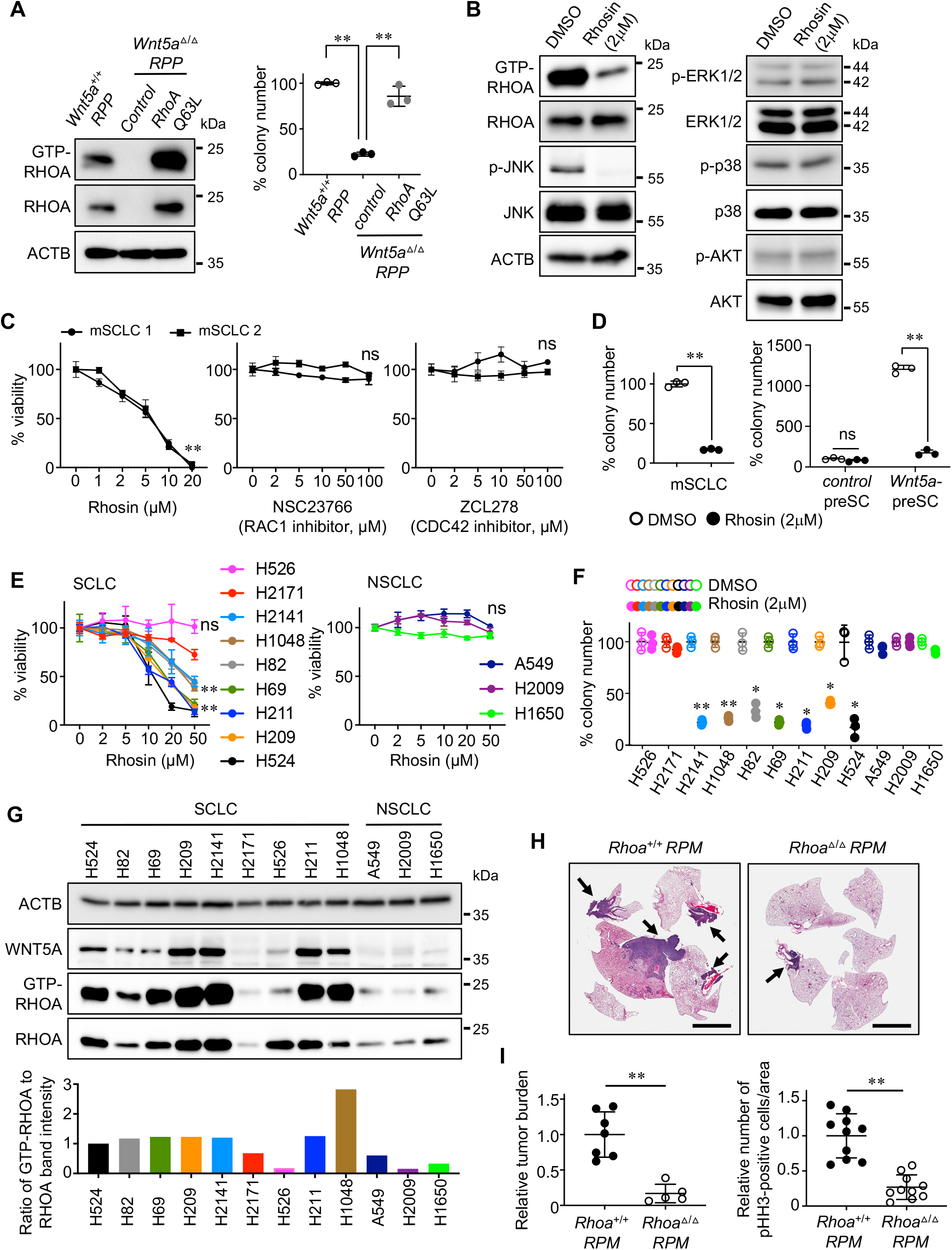
Activated RHOA is required for WNT5A-mediated SCLC development and cell proliferation. **(A)** Immunoblot for total and GTP-bound RHOA in *Wnt5a*^*+/+*^ and *Wnt5a*^*Δ/Δ*^ *RPP* cells, as well as *Wnt5a*^*Δ/Δ*^ *RPP* cells engineered to express a dominant-negative form of RHOA (RHOA Q63L) (left). Quantification of soft agar colonies (>0.2 mm in diameter) derived from these cells (n=3 replicates per cell type) (right). **(B)** Immunoblot for WNT5A, total and GTP-bound RHOA, and total and phosphorylated substrates of RHOA (JNK, ERK1/2, p38, and AKT) in mSCLC cells treated with Rhosin, a selective inhibitor of RHOA. **(C)** Results of MTT cell viability assays of 2 mSCLC cell lines treated with Rhosin and selective inhibitors of RAC1 (NSC23766), and CDC42 (ZCL278) (n=3 per cell type). **(D)** Quantification of soft agar colonies (>0.2 mm in diameter) formed from mSCLC cells or from control and *Wnt5a*-preSC cells treated with Rhosin or DMSO (n=3 per cell type). **(E)** Results of MTT cell viability assays of human SCLC and NSCLC cells treated with Rhosin (n=3 per cell type). **(F)** Quantification of soft agar colonies (>0.2mm in diameter) formed from human SCLC cells treated with Rhosin or DMSO throughout the assay (n=3 per cell type). **(G)** Immunoblot for WNT5A and total and GTP-bound RHOA in human SCLC and NSCLC cell lines. Levels of GTP-bound RHOA relative to total RHOA were calculated from band intensities and displayed below. ACTB was used as a loading control. **(H)** Representative images of H&E-stained lung sections and quantification of tumor burden (tumor area/lung area) in the lungs of *Rhoa*^*+/+*^ vs *Rhoa*^*Δ/Δ*^ *RPM* (*Rb/p53/Myc*^*T58A*^) mice (n=7 and n=5, respectively). Arrows indicate tumors. **(I)** Representative images of immunostained lung sections and quantification of pHH3-positive cells per lung area in *Rhoa*^*+/+*^ *vs. Rhoa*^*Δ/Δ*^ *RPM* (n=10 tumors per genotype). *, *p*<0.01; **, *p*<0.001. Statistical tests were performed using unpaired t-test (ns: not significant). Error bar represents standard deviation. Scale bar: H, 5 mm.

To determine the necessity of RHOA for SCLC development *in vivo*, we conditionally deleted *Rhoa* in the lungs of *RPM* mice. *RPM* mice were shown to develop tumor that lacks p130 and expresses WTN5A and RHOA (Fig. 3E and Supplementary Fig. 7). Eight weeks after intratracheal instillation of adenovirus CGRP-Cre, *Rhoa*^Δ/Δ^ *RPM* mice had significantly less tumor burden in the lungs than *Rhoa*^+/+^ *RPM* mice, while lung tumors in both genotypes displayed histology typical of SCLC (Fig. 7H, I and Supplementary Fig. 8). Quantification of pHH3-positive cells showed significantly fewer mitotic cells in the lung tumors of *Rhoa*^Δ/Δ^ vs *Rhoa*^+/+^ *RPM* mice (Fig. 7I and Supplementary Fig. 8). Taken together, these data suggest that RHOA is critical for SCLC development and cell proliferation.

## Discussion

The goals of this study were to define the importance of WNT signaling in SCLC and identify the molecular players involved. Our results show that beta-catenin is not required for SCLC development and cell proliferation despite reporter activity showing expression of its transcriptional target, *Axin2*, in SCLC cells. Instead, WNT5A was determined to drive the Axin2 reporter activity observed, promote the neoplastic transformation of precancerous precursor cells, and be required for SCLC development and cell proliferation. Furthermore, our data point toward a novel mechanism in which loss of p130 induces *Wnt5a* expression and WNT5A selectively induces and activates RHOA among the Rho family of GTPases to promote cell proliferation and survival. This WNT5A-RHOA axis appears specific to SCLC and may thus reveal a novel vulnerability for pharmacological intervention.

The dispensability of beta-catenin for SCLC is remarkable, given the canonical role of the WNT/beta-catenin pathway in oncogenic signaling. Our findings contribute to a growing body of evidence that the WNT/beta-catenin pathway is not required for the development of several cancer types, including malignant fibrous histocytoma, melanoma, and osteosarcoma (*13, 15, 16*). Further, ectopic beta-catenin has been demonstrated to inhibit cell proliferation, leading several studies to suggest that its tumor suppressor role is lost in some tumor types; notably in these cases, WNT5A was shown to promote tumor development by antagonizing the WNT/beta-catenin pathway (*14, 15*). Our studies in autochthonous mouse models and human cell lines strongly suggest that the beta-catenin-dependent pathway is neither oncogenic nor tumor-suppressive in SCLC. These data are consistent with our observation that biomarkers for WNT/beta-catenin pathway activity such as MYC and CCND1 are not differentially expressed between precancerous cells and tumor cells or upon beta-catenin loss. However, given emerging evidence on the plasticity of SCLC cells, we do not rule out the possibility that the WNT/beta-catenin pathway may promote SCLC cell survival in certain contexts, as may be the case in relapse following chemotherapy (*33*).

The molecular origin of WNT5A adds new insight into the increasing heterogeneity of SCLC that has recently begun to emerge. Molecular SCLC subtypes (SCLC-A/N/P) were proposed based on the expression patterns of ASCL1, NEUROD1, and POU2F3, and linked to different sensitivities to platinum-based, immune-based, and targeted therapies (*51*-*53*). Because the SCLC cell lines used in this study include SCLC-A/N/P subtypes, our findings suggest that WTN5A signaling may be implicated broadly in SCLC. Additionally, our findings show that *p130* knockout and restoration were sufficient to induce and suppress WNT5A expression, respectively, identifying p130 as an upstream regulator for WNT5A. These findings are notable in light of our discovery of RHOA as a downstream effector, as the frequent loss of p130 function in SCLC, while not actionable in and of itself (*5*), may serve as a biomarker for therapeutic approaches targeting the WNT5A-RHOA axis.

The transduction of WNT5A signaling via RHOA provides new insight into the complexity of WNT pathways. Previous studies suggested that WNT5A signaling via the WNT/PCP and WNT/Ca^2+^ pathways promotes cell invasion and chemoresistance in melanoma, mainly by driving cytoskeletal rearrangement and the epithelial-mesenchymal transition (EMT) (*54-57*). Other studies suggested that WNT5A suppresses colorectal cancer and lymphoid malignancies (*55, 56, 58-60*). These heterogeneous outcomes of WNT5A signaling may stem from different combinations of ligands, receptors, and transducers being present and active in a cell type-specific manner (*61*). Our findings suggest that the role of WNT5A in SCLC lies primarily in driving cell proliferation and survival, although its role in cytoskeletal rearrangement and EMT remains to be tested. Our data also point to RHOA as a key player in WNT5A-driven cell proliferation, although the exact mechanism of channeling WNT5A signaling into enhanced proliferation remains unknown. Based on previous studies, we surmise that increased RHOA activity in SCLC cells may accelerate the G1-S transition while also facilitating mitosis (*62-66*). Our findings suggest that this process appears to specifically involve activation of JNK instead of other well-known effectors, including ROCK1, p38, ERK1/2, RAC1, and CDC42, and does not rely on the phosphorylation of myosin light chain (MLC2) critical for cytoskeletal rearrangement. Thus, the WNT5A-RHOA axis appears to be not only specific to SCLC, but also distinct from other noncanonical WNT pathways such as the WNT/PCP and WNT/Ca^2+^ pathways.

The link between WNT5A induction and RHOA activity is notable and merits further study, given that it does not extend to the paralogs RHOB and RHOC, or to RAC1 and CDC42. One major mechanism of regulating Rho family members involves the guanine nucleotide exchange factors (GEFs) and GTPase-activating proteins (GAPs), which promote switching between the GTP-bound active form and GDP-bound inactive form of Rho family members (*67-71*). We do not rule out the possibility that WNT5A plays a role in this process to regulate RHOA post-transcriptionally, especially given the correlation between protein levels of WNT5A and GTP-bound RHOA. Although current understanding of these mechanisms is limited, our data nevertheless establish RHOA expression and activity as robust biomarkers for WNT5A pathway activation in SCLC.

Another question that remain to be addressed is how WNT5A influences *Axin2* expression and what role AXIN2 plays in SCLC. Our gene expression analyses and use of the *Axin2*-*lacZ* reporter suggest that *Axin2* is a downstream target of the WNT5A pathway, but are not sufficient to demonstrate a link between the WNT5A-AXIN2 and WNT5A-RHOA axes. Likewise, this study does not focus on potential oncogenic roles for other WNT ligands involved in beta-catenin-independent pathways, even though we found enrichment of genes encoding these ligands in SCLC relative to normal lung tissue. While these WNTs remain poorly characterized in SCLC, a recent study has linked WNT11 activation to the growth of ASCL1-positive SCLC cells (*34*). We therefore surmise that SCLC heterogeneity may include multiple beta-catenin-independent pathways playing a role in tumor development.

This study implicates a novel, WNT5A-driven pathway in the development and continuing growth of SCLC and provides critical insight into the development of therapeutic strategies, as well as the stratification of patients who may benefit from them. With development of a clinically applicable RHOA inhibitor underway, future experiments can focus on further assessing the scope and impact of inhibiting the WNT5A-RHOA axis on human SCLC.

## Materials and Methods

### Mouse strains, tumor induction, and allografts

Strains carrying *p53*^*lox*^, *Rb*^*lox*^, *p130*^*lox*^, *Axin2*^*lacZ*^, *beta-catenin*^*lox*^, *Rhoa*^*lox*^, and *H11*^*lox-stop-lox-MycT58A*^ alleles were generously shared by Drs. Anton Berns, Tyler Jacks, Julien Sage, Walter Birchmeir, Rolfk Kemler, Trudy Oliver and Robert Wechsler-Reya, respectively (*40, 42, 48, 72-76*). Generation of compound transgenic mice, including *Rb*^*lox/lox*^ *p53*^*lox/lox*^ *p130*^*lox/lox*^ (*RPP*) mice and *Rb1*^*lox/lox*^ *Trp53*^*lox/lox*^ *H11*^*lox-STOP-lox* -*MycT58A*^ (*RPM*) mice, has been previously described (*42, 48*). All mice were genotyped before and after experiments using polymerase chain reaction (PCR) for tail DNA. DNA was purified using lysis buffer containing Proteinase K (Thermo Fisher Scientific, BP1700-100) and genotyping PCR was performed using the primers described in Supplementary Table 1. For tumor induction, lungs of 10-week-old mice were infected with adenoviral Cre (Ad-Cre) via intratracheal instillation as previously described (*77*) and mice were aged 6 months. Multiple cohorts of independent litters were analyzed to control for background effects, and both male and female mice were used. Ad-Cre was purchased from the University of Iowa Gene Transfer Vector Core. For allograft experiments, 5.0 × 10^5^ human or murine cells were injected in the flanks of nude mice (*Foxn1*^*nu/nu*^, Envigo). Following implantation, mice were fed a doxycycline diet (625 mg/kg, Envigo). Perpendicular tumor diameters were measured using calipers. Volume was calculated using the formula L × W^2^ × 0.52, where L is the longest dimension and W is the dimension perpendicular to L. Injected mice were maintained and observed for palpable tumors according to procedures approved by IACUC and euthanized when tumor size reached 1.5 cm in diameter, the endpoint of allograft study per institutional animal policy.

### Cells, proliferation and soft agar assay

293T cells were cultured in normal DMEM media supplemented with 10% bovine growth serum (GE Healthcare, SH30541.03) and 1% penicillin-streptomycin (Invitrogen, 15140-122). Mouse SCLC cells and precancerous cells (preSC) were derived from lung tumors and early-stage neuroendocrine lesions, respectively, that developed in the *Rb1/Trp53-*mutant model (*36, 42*)). Human SCLC cell lines (NCI-H69, NCI-H82, NCI-H209, NCI-H524, NCI-H2141, NCI-H2171, NCI-H211, NCI-H526, NCI-H1048, NCI-A549, NCI-H1650, NCI-H2009) were gifts of Adi Gazdar and John Minna (UT Southwestern), Julien Sage (Stanford), Hisashi Harada (Virginia Commonwealth University), and Christopher Vakoc (Cold Spring Harbor Laboratory). These cell lines were authenticated by profiling patterns of seventeen short tandem repeats (ATCC, 135-XV and 200FTA), and tested negative for mycoplasma using the PlasmoTest-Mycoplasma Detection Kit (InvivoGen, rep-pt1). Both human and mouse cells were cultured in RPMI1640 media supplemented with 10% BGS and 1% penicillin-streptomycin. For screening the WNT pathway ligands, mouse SCLC cells were seeded into 6-well plates and cultured for 16 h in RPMI1640 + 10% BGS with or without 100 ng/mL of Wnt5a (R&D System, 645-WN-010) or recombinant Wnt3a (PeproTech, 315-20) in an incubator at 37 °C, 5% CO_2_, 3% O_2_. For soft agar assay to measure clonal expansion, cells were suspended at 1×10^4^ cells per well in 0.5 mL of growth medium containing 0.35% low-melting-point agarose (Invitrogen, 16520-100), and seeded on top of a 0.5 mL base layer of medium containing 0.5% agar. Cells were allowed to grow at 37°C with 5% CO_2_ for 3 weeks, and medium was changed every 3 days. Colonies were fixed with 10% MeOH and 10% acetic acid at room temperature for 10 minutes, then stained with 1% MeOH, 1% Formaldehyde, and 0.05% crystal violet. Low-melting-point agarose was premixed with RPMI 2X (Fisher Scientific, SLM202B) complemented with 20% BGS, 200 Unit/mL penicillin, and 200 μg/mL streptomycin. Images of wells are acquired using the Olympus MVX10 scope, and colonies from whole-field images were counted using NIS-Elements Basic Research (Nikon) imaging software. All cell culture experiments were performed in triplicates and repeated for a minimum of two biological replicates.

### Chemical, vectors and virus production

Lentiviral plasmid pCW57.1 (a gift from David Root) was used to express murine *Wnt5a* cDNA. Tet-pLKO-puro (a gift from Dmitri Wiederschain) was used to express short hairpin RNAs (shRNAs) against human *WNT5A*. pL-CRISPR.EFS.tRFP (a gift from Benjamin Ebert) and LV-gRNA-Zeocin (a gift from Mazhar Adli) were used to express Cas9 and guide RNA (gRNA) against murine targets. *Wnt5a* cDNA was cloned using total RNAs from murine SCLC cells and its sequence and protein expression were verified. All shRNA and gRNA sequences are included in Supplementary Table S1. Lentivirus was produced by co-transfecting lentiviral plasmids with packaging plasmids, psPAX2 and pMD2.G (gifts from Didier Trono), into 293T cells using polyethylenimine (Sigma-Aldrich, 408727). For lentiviral infection, supernatants containing the viral particles were harvested 48 and 72 hours after transfection of 293T cells, filtered through 0.45μm PVDF filters, and added to cell culture in the presence of 5μg/ml polybrene (Sigma-Aldrich, H9268). To generate KO cells, parental cells were infected with the LV-gRNA-zeocin vector carrying gene-specific or non-targeted (control) sgRNAs, using Lipofectamine 2000 (Invitrogen, 52887) according to manufacturer instructions. Forty-eight hours later, selection of infected cells was performed using zeocin (Thermo Fisher Scientific, R25001) or puromycin (Thermo Fisher Scientific, A1113803). Sanger sequencing and immunoblot were used to verify mutations in targeted sequences and loss of targeted proteins, respectively. Inhibitors of the RHOA subfamily (Rhosin, 555460), RAC1 (NSC23766, 553502), and CDC42 (ZCL278, 5.00503) were purchased from Sigma-Aldrich. Retrovirus was produced by co-transfecting retroviral plasmids with packaging plasmids, VSV-G and gag-pol, into 293T cells using polyethylenimine. For retroviral infection, supernatants containing the viral particles were harvested 48 and 72 hours after transfection of 293T cells, filtered through 0.45μm PVDF filters, and added to cell culture in the presence of 5μg/ml polybrene. To introduce a constitutively active form of RHOA, cells were infected with empty (control) MIEG3 vector or vector carrying the RHOA Q63L sequence, and GFP-positive cells were sorted 48 hours later using FACS (BD Influx).

### Histology, immunostaining, immunoblot, and X-gal staining

Lungs were perfused with and incubated in 4% paraformaldehyde (PFA)/phosphate-buffered saline (PBS) overnight before paraffin embedding. Five-micron (5µm) sections were used for hematoxylin and eosin (H&E) staining and immunostaining. To perform immunostaining, sections were de-waxed and hydrated using Trilogy (Cell Marque). The primary antibodies used are listed in Supplementary Table 3. Alexa Fluor-conjugated secondary antibodies (Invitrogen) were used for visualization, and anti-fade reagents with DAPI (Vector Lab) were used to preserve fluorescence and stain for nuclei. For quantification of phosphorylated histone H3 (pHH3)-positive cells and CGRP staining, tumors of similar size and area were included. Macroscopic images of H&E-stained lung sections, and microscopic images of H&E-stained and immunostained tissues, were acquired using Olympus MVX10 (Olympus) and Nikon Eclipse Ni-U microscope (Nikon), respectively. Image analysis and automated quantification were performed using NIS-Elements Basic Research (Nikon). To perform immunoblot analysis, cells were lysed in RIPA buffer (50 mM Tris-HCl pH7.4, 150 mM NaCl, 2 mM EDTA, 1% NP-40, 0.1% SDS) and sonicated for 20s at 15% intensity. Generally, 15∼30 µg protein lysate was loaded into various percentages of SDS-polyacrylamide gels for electrophoresis in PAGE running buffer (25 mM Tris, 192 mM glycine, 0.1% SDS, pH8.3). Proteins were then transferred to 0.2 or 0.45 μm PVDF membrane in transfer buffer (25 mM Tris, 192 mM glycine, 20% methanol). Membranes were blocked for 1 hour at room temperature using TBST buffer (150 mM NaCl, 10 mM Tris pH8.0, 0.1% Tween20) supplemented with 5% non-fat milk, then blotted with primary antibodies overnight at 4°C on an orbital shaker. After washing 5 times with TBST, membranes were incubated with secondary antibodies for 2 hours at room temperature and washed another 5 times with TBST. Membranes were then incubated with ECL Western Blotting Detection Reagent (Thermo Fisher Scientific, 32106) for 5 minutes, and signals were detected using a ChemiDoc machine (Bio Rad). All primary and secondary antibodies used are listed in Supplementary Table 3. To perform X-gal staining, lungs were inflated in 4% PFA/PBS for 10 minutes, washed in 0.02% NP-40/PBS for 10 minutes, and incubated in 1mg/mL X-gal in 0.02% NP-40/PBS overnight at room temperature. X-gal-stained lungs were washed twice in 0.02% NP-40/PBS and fixed in 4% PFA/PBS for 24 hours before whole-mount imaging and paraffin embedding.

### Assays for activation of Rho family GTPases and ROCK kinase

Rho activation assays for RHOA, RAC1, and CDC42 were performed using Small GTPase Activation Kits according to the manufacturer’s protocol (Cell Biolabs; STA-401, 402, 403). Briefly, cells at 70%–80% confluence were lysed for 15 minutes on ice, lysate was centrifuged for 10 minutes at 14,000g and 4°C, and supernatant was then incubated with 40 μL of Rho-binding domain (RBD) beads for 1 hour on ice. Subsequently, the beads were centrifuged for 20 seconds at 14,000g and 4°C, supernatant was removed, and 0.5 ml of lysis buffer was added to the beads; this step was repeated three times. An equal volume (40 μL) of 2× reducing SDS-PAGE sample buffer was then added to the bead solution, which was then boiled at 95°C for 10 minutes. After centrifugation for 30 seconds at 14,000g and 4°C, 20 μL of the sample was used for immunoblotting with anti-RHOA/RAC1/CDC42 antibodies included in the assay kits. The assay for ROCK1 kinase activity assay was performed according to the manufacturer’s protocol (Cell Biolabs, STA-415). Briefly, cells were washed twice with ice-cold PBS and lysed with an immunoprecipitation buffer (50 mm Tris-HCl, pH 7.5, 150 mM NaCl, 1 mM PMSF, 1 mM EDTA, 1% Triton X-100, and mixture proteinase inhibitor). Cell lysates were centrifuged at 13,000 rpm for 10 min to remove insoluble debris. Supernatants were transferred to fresh tubes, incubated with an anti-ROCK1 antibody for overnight at 4 °C. Protein A/G agarose beads (GenDEPOT, P9203) were then added for 4 h with agitation at 4 °C. The beads were washed three times in immunoprecipitation buffer and incubated with the kinase reaction mixture, following the protocol indicated by the manufacturer. We determined phosphorylation of the endogenous ROCK1 substrate MYPT1 in immunoprecipitates using anti-phospho-Thr696-MYPT1 antibody.

### Quantitative RT-PCR (RT-qPCR) and RNA sequencing

For RT-qPCR, total RNA was isolated from cells or tumors using TRIzol (Life Technologies) according to manufacturer protocol. cDNA was generated by the ProtoScript II first strand cDNA synthesis kit (New England Biolabs, E6560). Quantitative PCR (RT-qPCR) was performed using SYBR Green (Thermo Fisher Scientific, 4367659) with Applied Biosystems 7900 or StepOnePlusTM following manufacturer protocol. All qPCR primer sequences were retrieved from the online Universal Probe Library at Roche and included in Supplementary Table 3. For RNA-seq analysis, total RNA was extracted using the RNEasy kit (Qiagen, 74106). Sequencing libraries were generated by the Genome Analysis and Technology Core at the University of Virginia using oligo dT-purified mRNA from 500ng of total RNA and the NEB Next Ultra RNA library preparation kit (New England Biolabs), and 50-bp single-end sequencing was performed on the Illumina NextSeq500 platform (Illumina). The sequencing data have been deposited to NCBI GEO Datasets (accession number GSE197893). Reads were mapped to the Mus musculus genome assembly GRCm38 (mm10) using TopHat and reads mapping to each gene were quantified using HTSeq (*78, 79*). Differentially expressed genes regulated by WNT5A were identified using DESeq2 package (*80*). Gene set enrichment analysis (GSEA) was performed for Hallmark gene sets from MSigDB R package (version 7.2.1) (*81*).

### Statistical analysis

Statistical analyses and graphing were performed with GraphPad Prism 8.2. Results are presented as the mean ± standard deviation and evaluated for significance at *p* value < 0.05 using an unpaired Student two-tailed t-test or, for Kaplan-Meier curves of lung tumor-free survival, a log-rank test.

## Acknowledgments

We thank Drs. Anton Berns, Tyler Jacks, Julien Sage, Walter Birchmeir, Rolfk Kemler, Trudy Oliver and Robert Wechsler-Reya for sharing strains carrying *p53*^*lox*^, *Rb*^*lox*^, *p130*^*lox*^, *Axin2*^*lacZ*^, *beta-catenin*^*lox*^, *Rhoa*^*lox*^, and *H11*^*lox-stop-lox-MycT58A*^ alleles, respectively. We thank Jenny Hsu, Julien Sage, and Debadrita Bhattacharya for their comments on the manuscript. We also thank the Research Histology Core and the Genome Analysis and Technology Core at the University of Virginia Comprehensive Cancer Center (P30CA044579), and the Biostatistics and Bioinformatics Shared Resources at the H. Lee Moffitt Cancer Center & Research Institute (P30CA076292). K.P. was supported by NIH grants (R01CA194461, U01CA224293) and American Cancer Society Research Scholar Grant (RSG-15-066-01-TBG). JP was supported by NIH grants (R01CA193297, R03CA256207) and The Cancer Prevention and Research Institute of Texas (RP200315).

## Author contributions

K-B.K. and K-S.P. contributed to conceptualization, methodology, data analysis, manuscript writing and reviewing. D-W.K., Y.K., J.T., N.K., and Y.G., contributed to methodology and data analysis. B.F. contributed to resources. JP contributed to conceptualization, data analysis and manuscript reviewing. Y.Z. contributed to resources, data analysis, and manuscript reviewing.

## Data and materials availability

All data needed to evaluate the conclusions in the paper are present in the paper and/or the Supplementary Materials. The RNA sequencing data have been posited to NCBI GEO Datasets (accession number GSE197893).

## Supplementary Materials

**Supplementary Fig. 1.**
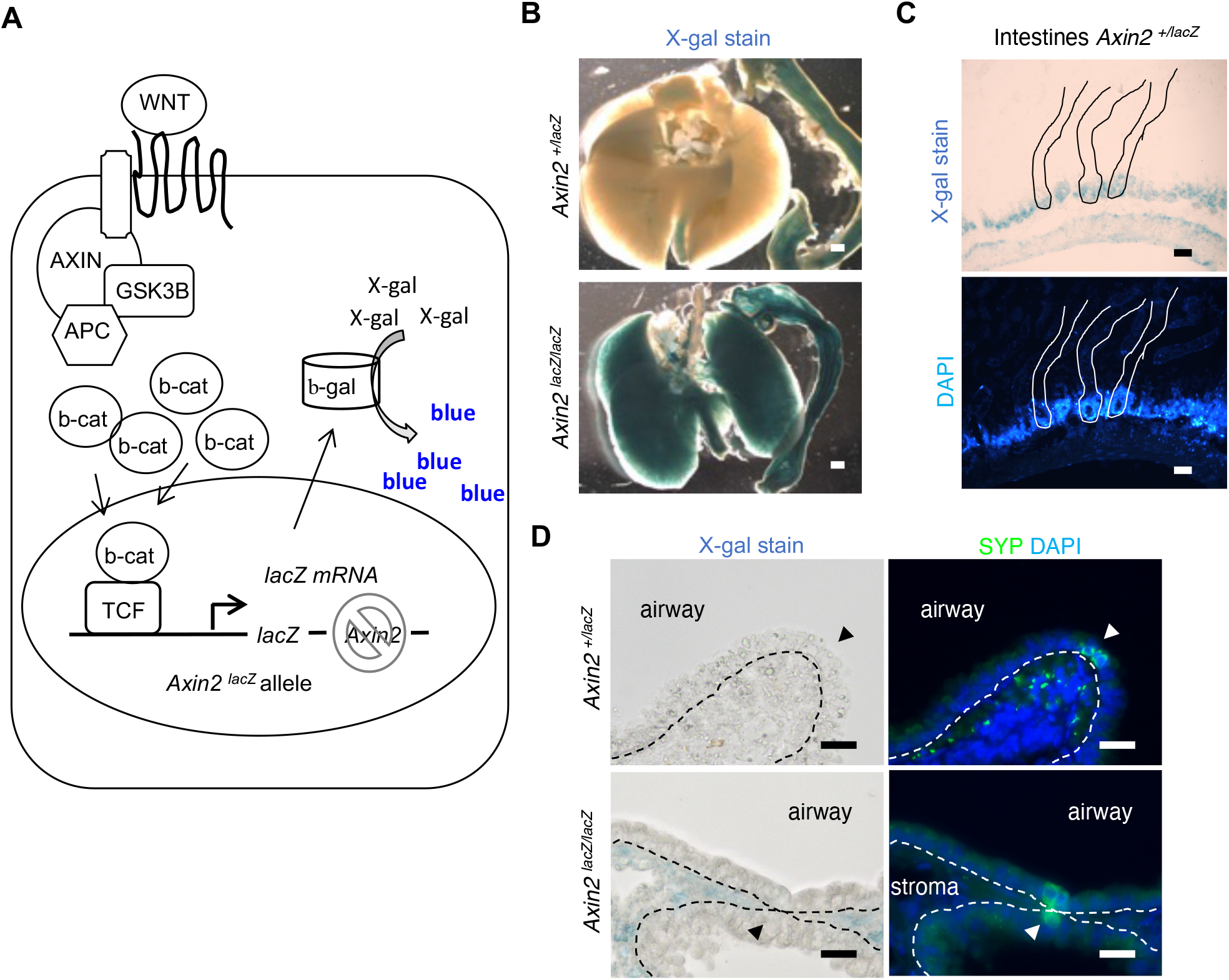
*Axin2-lacZ* reporter activity in *Axin2*^*+/lacZ*^ and *Axin2*^*lacZ/lacZ*^ mice. (A) Diagram of active WNT/beta-catenin pathway depicting major pathway components and the reporter gene *lacZ*, encoding beta-galactosidase (b-gal), knocked in the *Axin2* gene locus. Following transcription of *lacZ*, beta-galactosidase turns X-gal (a colorless cell-permeable substrate) into blue products accumulating in cells, which visualizes cell and tissues with the pathway activity. (B) Whole-mount images of X-gal-stained lungs and intestines in *Axin2*^*+/lacZ*^ and *Axin2*^*lacZ/lacZ*^ mice. Once induced by the pathway, AXIN2 forms a negative-feedback regulator of beta-catenin level. Therefore, loss of AXIN2 increases the WNT/beta-catenin (b-cat) pathway activity and constitutively activate the *Axin2* promoter that control the lacZ reporter expression, resulting in a drastic increase in X-gal staining as observed in the lung and intestine of *Axin2*^*lacZ/lacZ*^ mice. (C) Images of light microscopy of an intestine section showing X-gal staining in crypts (top) and fluorescence microscopy of the same section stained with DAPI (bottom). Lines demarcate individual crypts. Strong DAPI staining in the bottom of crypt indicate a high proliferation rate. (D) Light microscopy images of X-gal-stained lung sections from *Axin2*^*+/lacZ*^ and *Axin2*^*lacZ/lacZ*^ mice (left), and representative images of immunostaining for synaptophysin (SYP), a marker of neuroendocrine cells (right). DAPI was used as a counterstain. Dotted lines indicate the boundaries between the lung epithelium and the underlying stroma. The lacZ reporter activity is detected in subsets of stroma cells but undetectable in the airway epithelium of both *Axin2*^*+/lacZ*^ and *Axin2*^*lacZ/lacZ*^ mice, suggesting the absence of active WNT/beta-catenin pathway in the normal airway epithelium. Scale bars: A, 1 mm; B, 50 μm; C-E, 10 μm.

**Supplementary Fig. 2.**
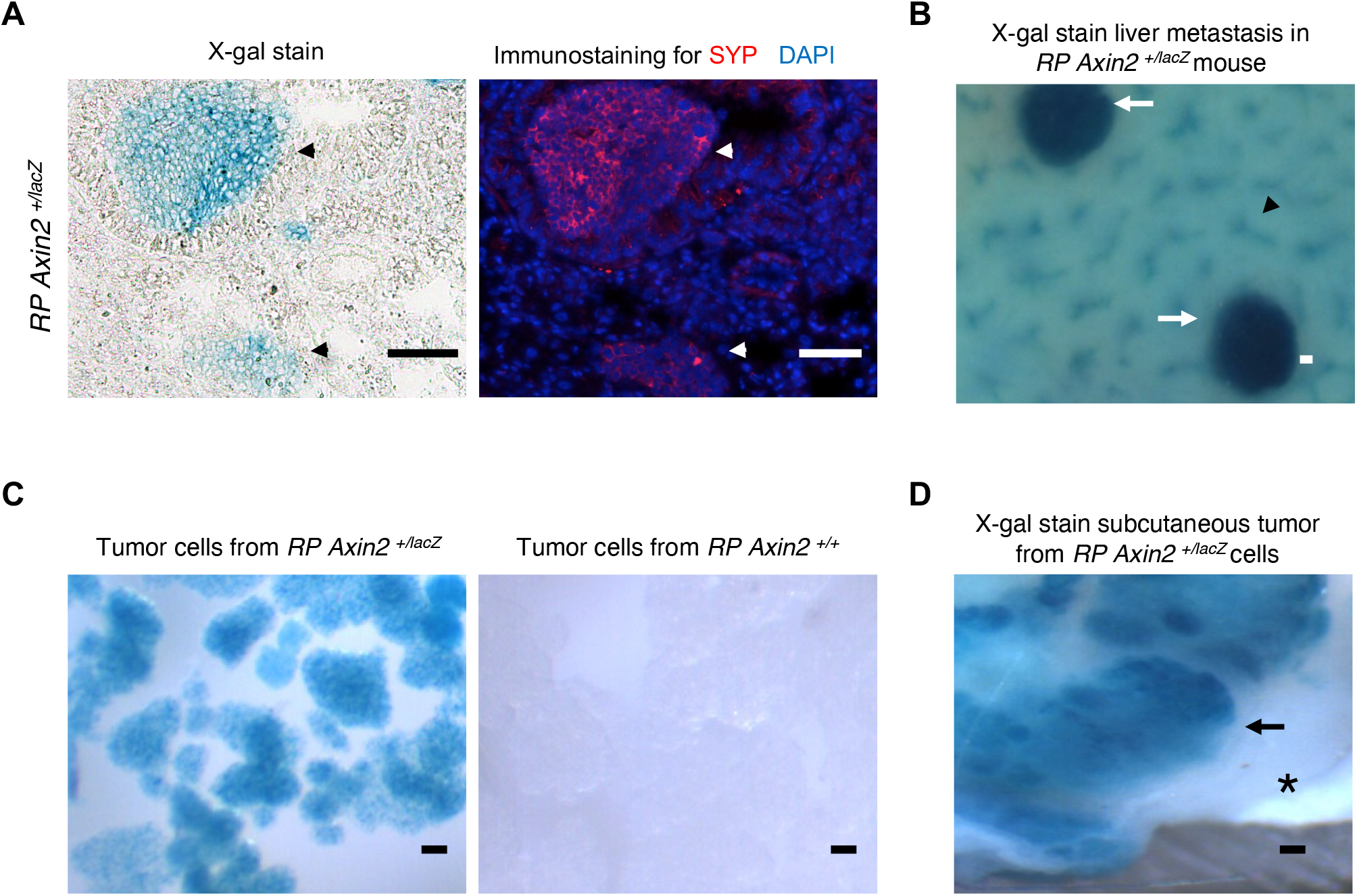
*Axin2-lacZ* reporter activity in mouse SCLC tumors and cells. (A) Representative images of light microscopy (left) and SYP immunostaining (right) on the same X-gal-stained lung section from *RP Axin2*^*+/lacZ*^ mice. DAPI was used as a counterstain. Arrowheads indicate small neuroendocrine lesions. (B) Image of whole-mount X-gal-stained liver from *RP Axin2+/lacZ* mice. Arrows indicate metastatic tumors, and arrowhead indicates normal portal triads. (C) Images of whole-mount X-gal-stained primary tumor cells derived from the lungs of *RP Axin2*^*+/lacZ*^ and *RP Axin2*^*+/+*^ mice. (D) Image of a whole-mount X-gal-stained subcutaneous tumor developed from *RP Axin2*^*+/lacZ*^ tumor cells. Arrow indicates tumor and asterisk indicates normal dermis of athymic nude mice. Scale bars: A and C, 50 μm; B and D, 0.1 mm.

**Supplementary Fig. 3.**
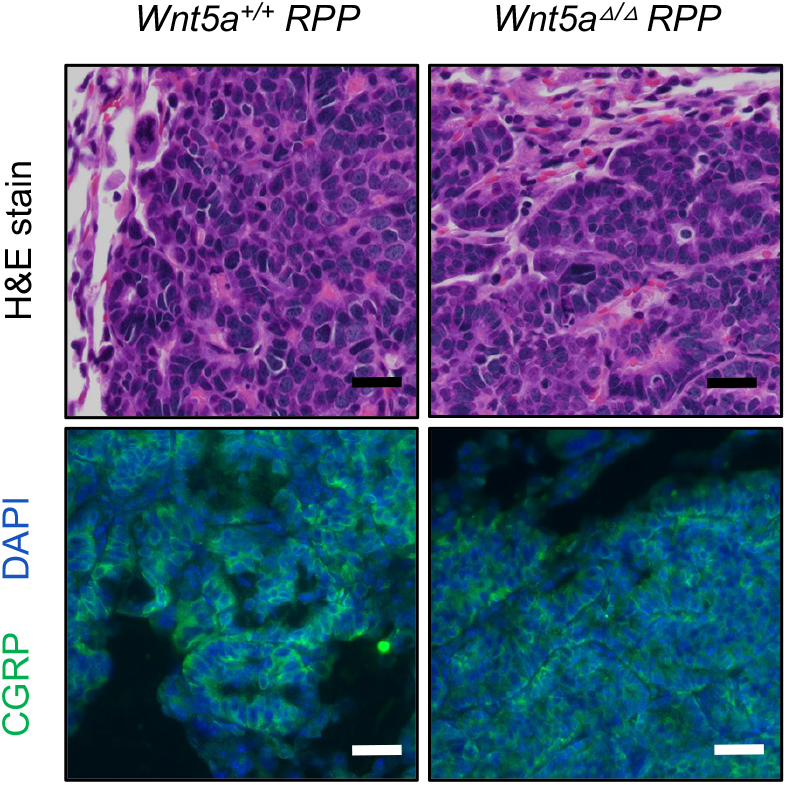
*Wnt5a* deletion does not affect the small cell histology and neuroendocrine differentiation of tumors developed in *RPP* mice. Representative images of H&E-stained lung sections of *Wnt5a*^*+/+*^ vs. *Wnt5a*^Δ/Δ^ *RPP* mice (top) and immunostaining for SYP (green, bottom). DAPI (blue) stains for nuclei. Scale bars: 50 μm.

**Supplementary Fig. 4.**
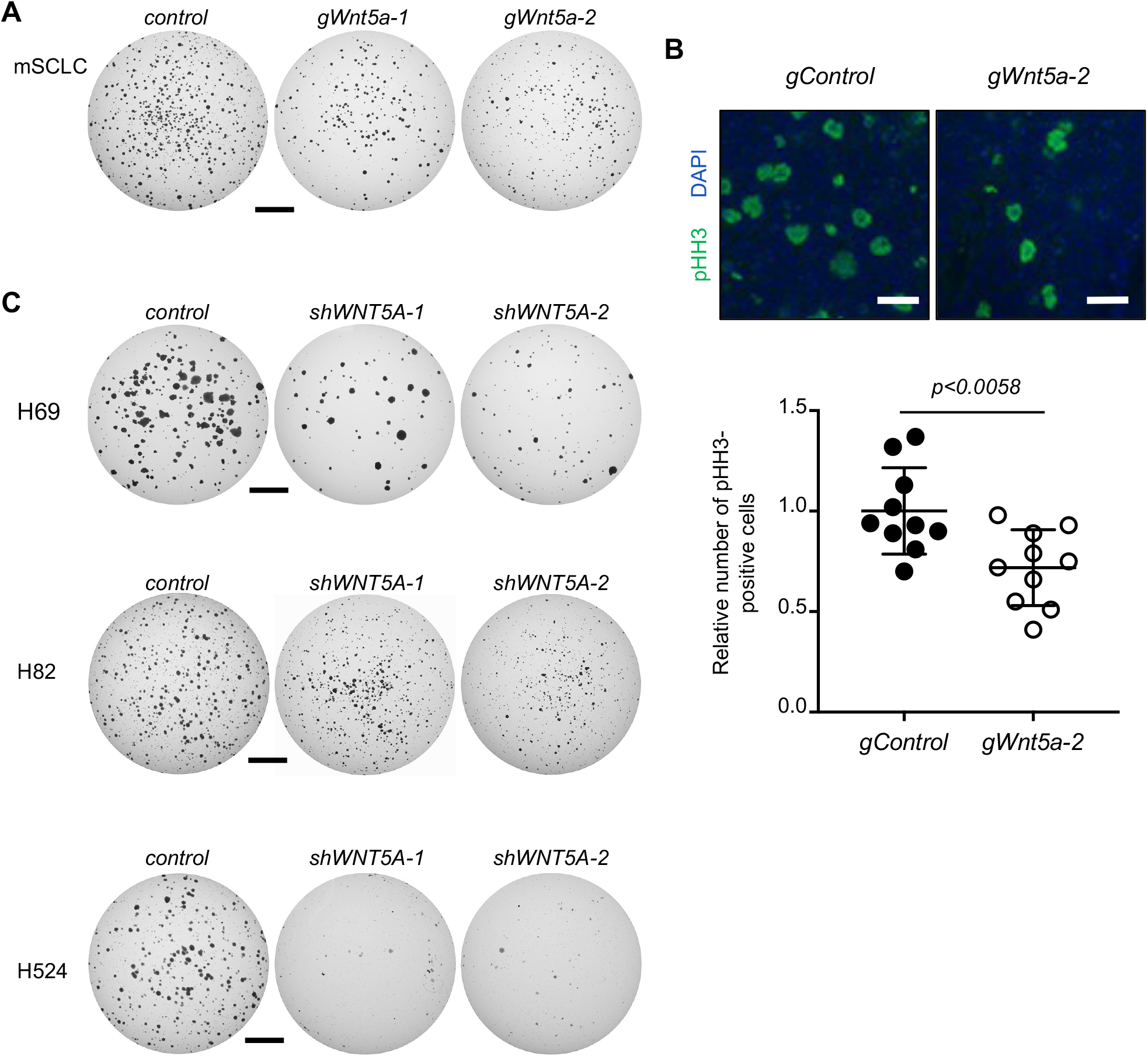
WNT5A loss suppresses the proliferation and tumorigenic potential of mouse and human SCLC cells. (A) Representative images of soft agar colonies derived from mouse SCLC (mSCLC) cells lacking *Rb1* and *Trp53*, treated with non-targeting gRNAs or gRNAs targeting *Wnt5a*. (B) Representative images of phospho-histone H3 (pHH3) immunostaining in sections of allograft tumors derived from the cells in A (top), and quantification of number of pHH3-positive cells. (C) Representative images of soft agar colonies derived from control or *WNT5A-*knockdown human SCLC cells. Statistical tests were performed using unpaired t-test. Error bars represent standard deviation. Scale bars: A, C, 5 mm; B: 50 μm.

**Supplementary Fig. 5.**
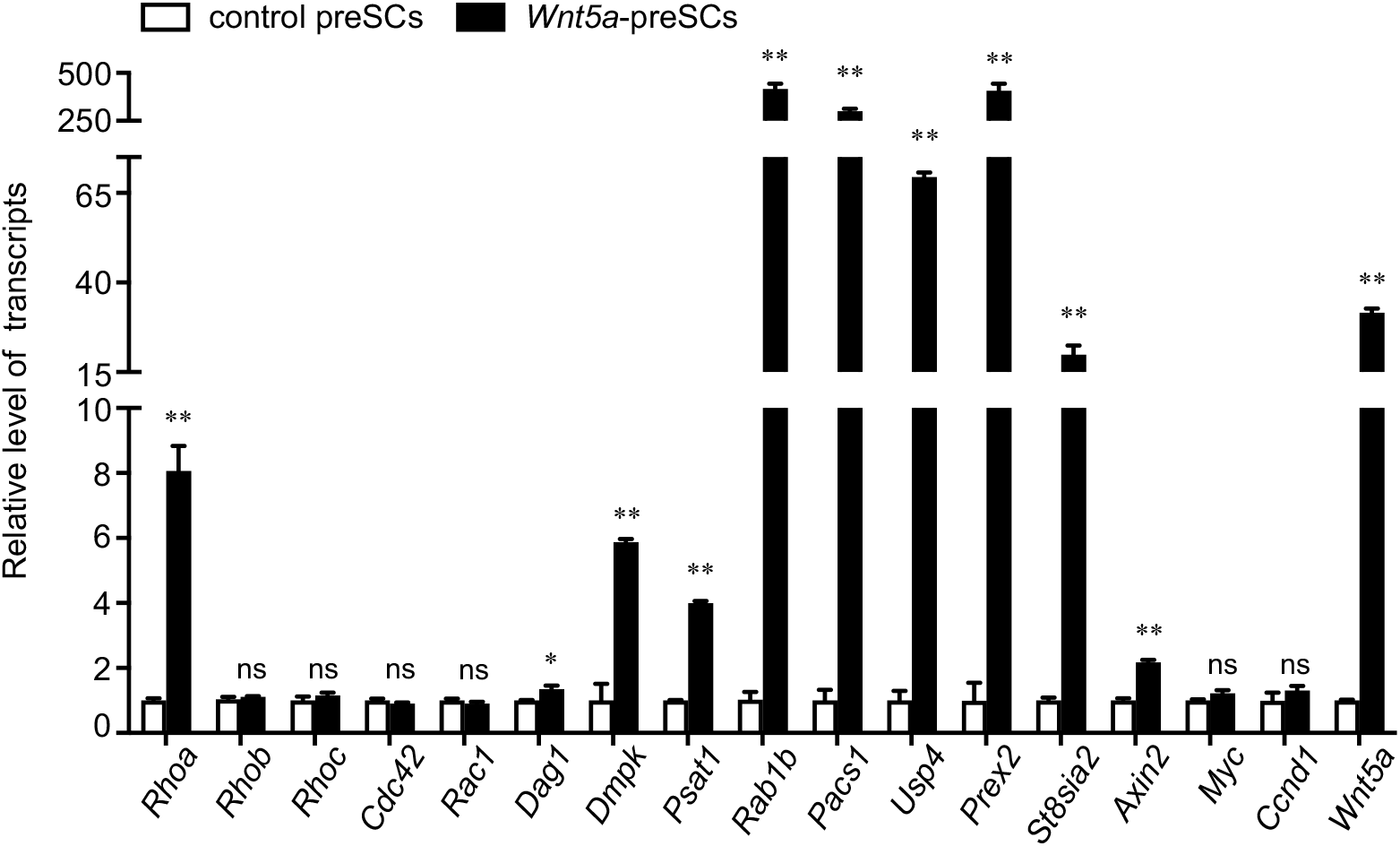
Validation of select genes differentially expressed in *Wnt5a*-preSCs. Plot of RT-qPCR showing a handful of DEs identified through RNA-seq in Fig. 6. *, p<0.05; **, p<0.01. Statistical tests were performed using unpaired t-test. Error bar represents standard deviation. ns: not significant.

**Supplementary Fig. 6.**
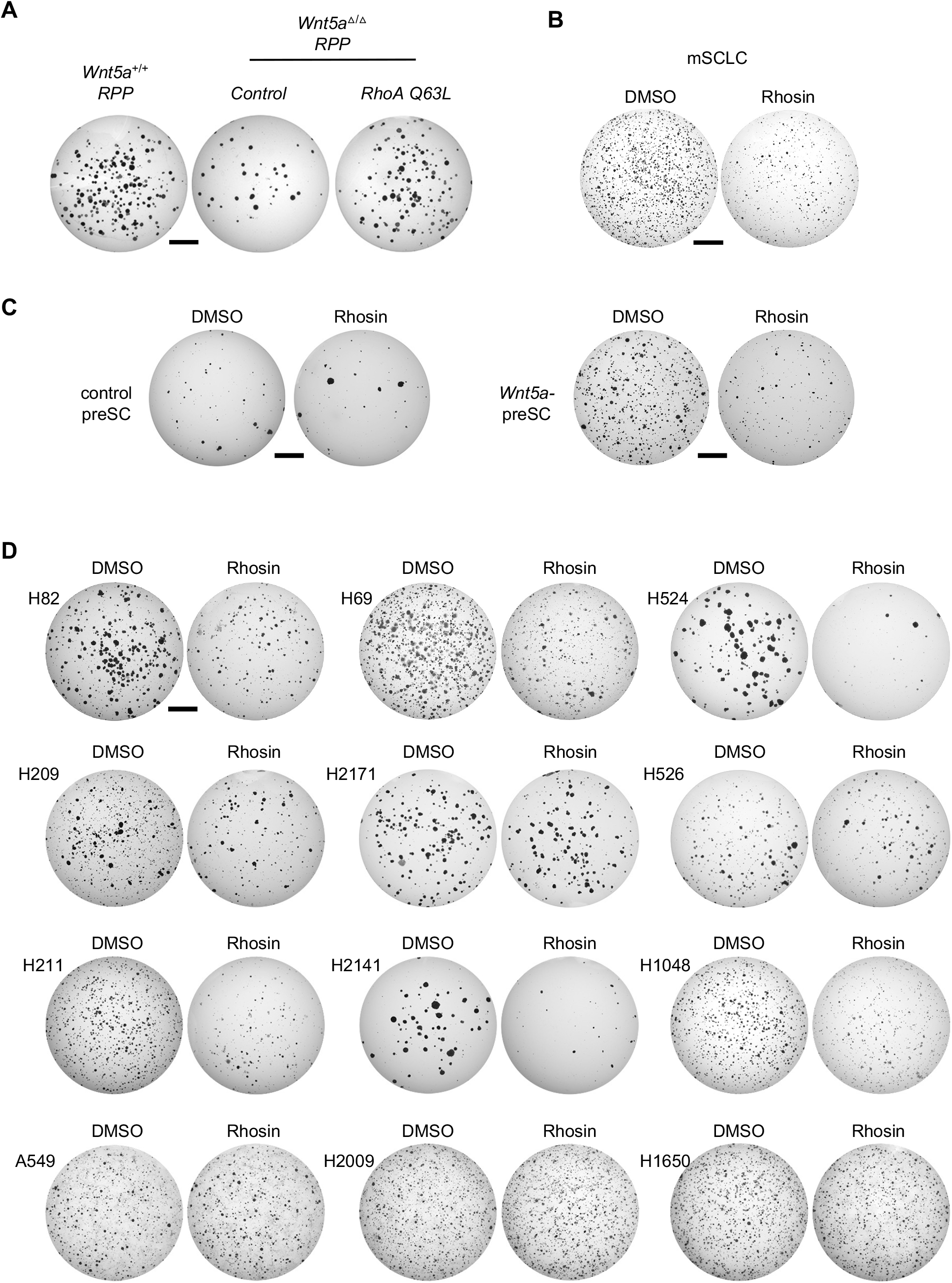
Inhibition of RHOA activity suppresses WNT5A-driven proliferation and tumorigenesis of mouse and human SCLC cells. (A) Representative images of soft agar colonies derived from *Wnt5a*^+/+^ *RPP* and *Wnt5a*^Δ/Δ^ *RPP* tumor cells as well as *Wnt5a*^Δ/Δ^ *RPP* tumor cells infected lentivirus expressing the dominant active form or RHOA (*RHOA Q63L*). (B) Representative images of soft agar colonies derived from mSCLC cells (*RP*) treated with DMSO or 2 μM Rhosin, a selective inhibitor of GTPase activity in the RHOA subfamily. (C, D) Representative images of soft agar colonies derived from control preSCs, *Wnt5a*-preSCs, and human SCLC lines treated with DMSO or 2 μM Rhosin. Scale bars: 5 mm.

**Supplementary Fig. 7.**
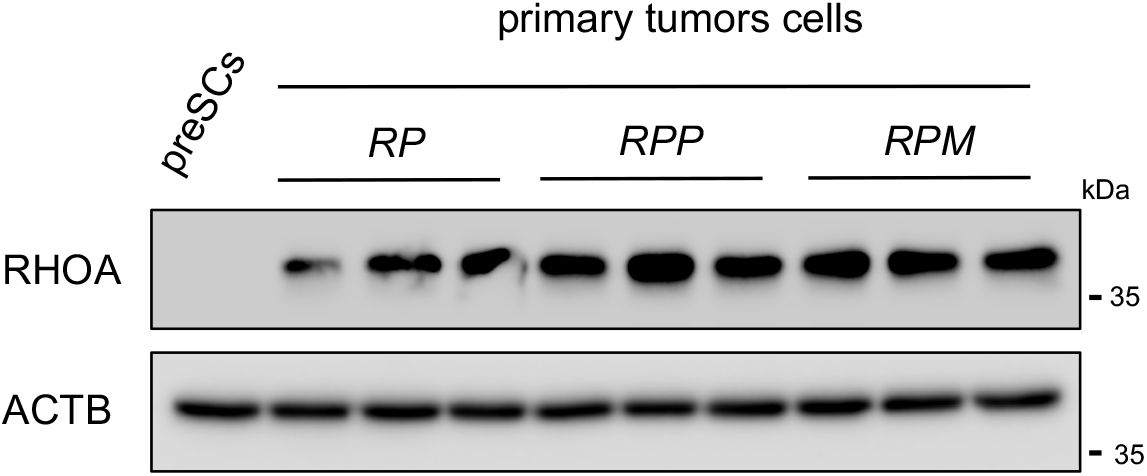
Increased RHOA expression in primary tumor cells derived from mouse SCLC tumors. Immunoblot shows RHOA proteins in primary cell derived from tumors developed in *RP, RPP*, and *RPM* mice, compared to the protein in preSCs. ACTB was used as a protein loading control.

**Supplementary Fig. 8.**
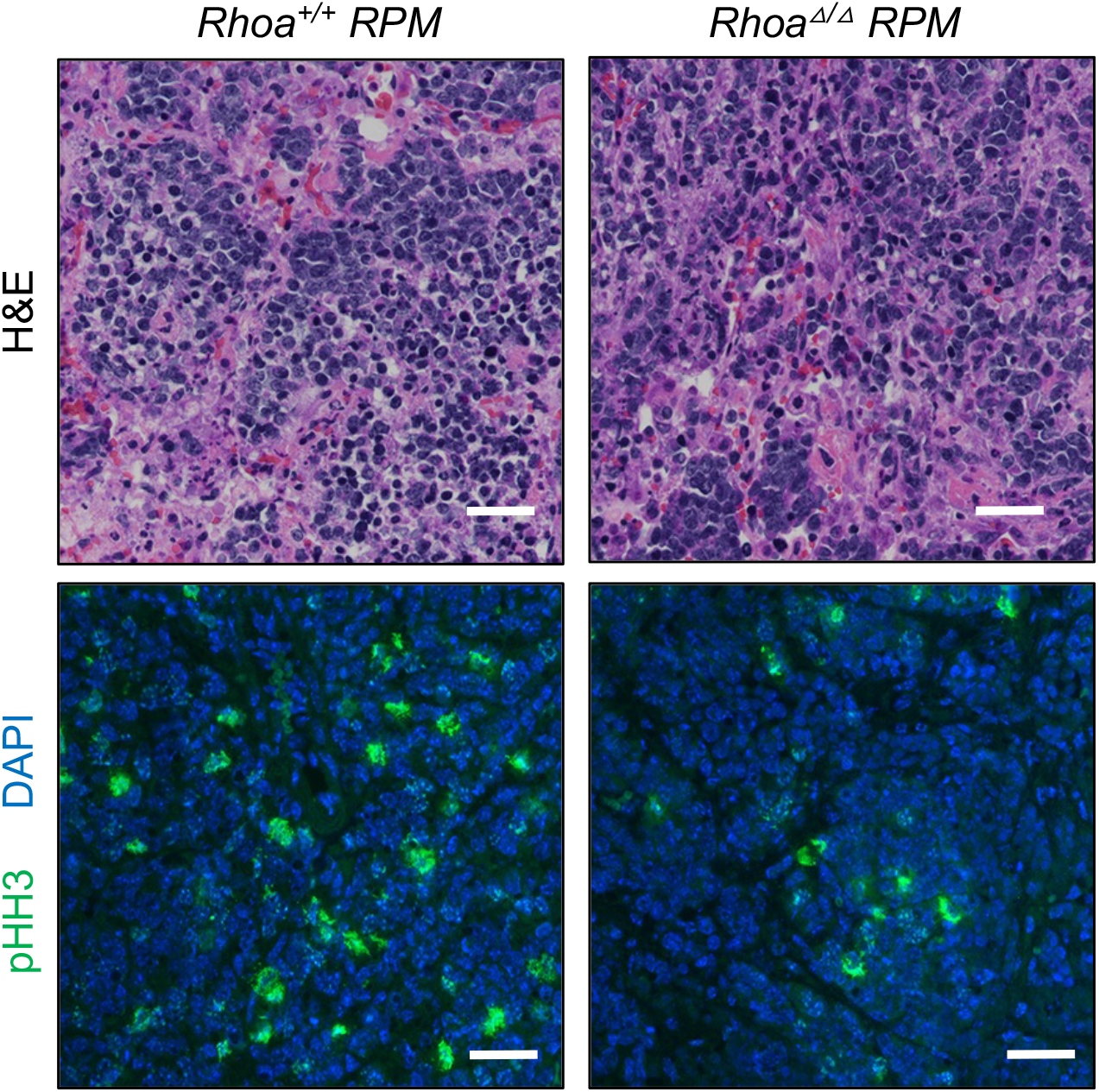
*Rhoa* deletion does not affect small cell histology but reduces cell proliferation in tumors developed in *RPM* mice. Representative images of H&E-stained lung sections of *Rhoa*^*+/+*^ vs. *Rho*^Δ/Δ^ *RPM* mice (top) and immunostaining for pHH3 (bottom). DAPI (blue) stains for nuclei. Scale bars: 50 μm.

**Supplementary Table 3.**
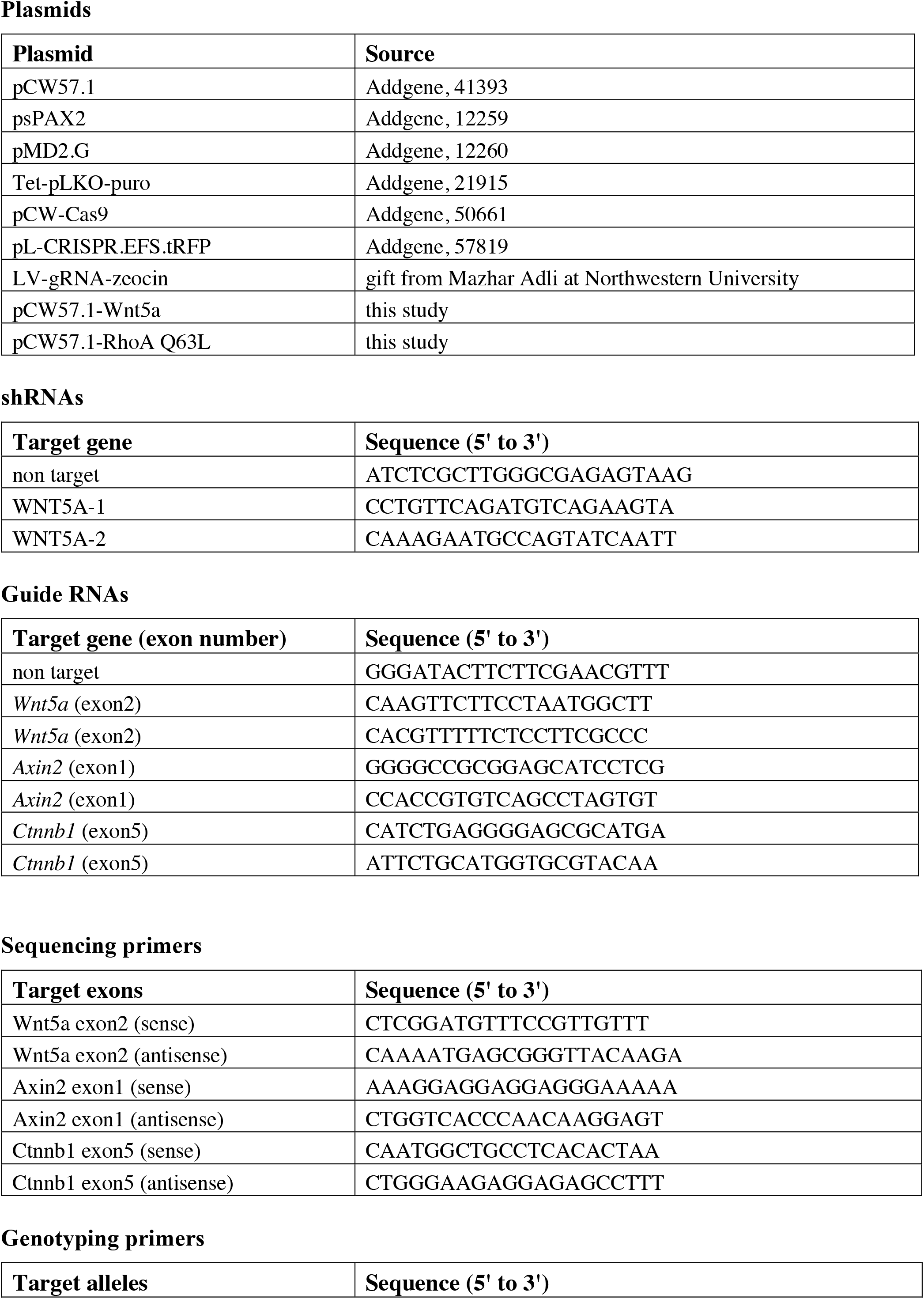
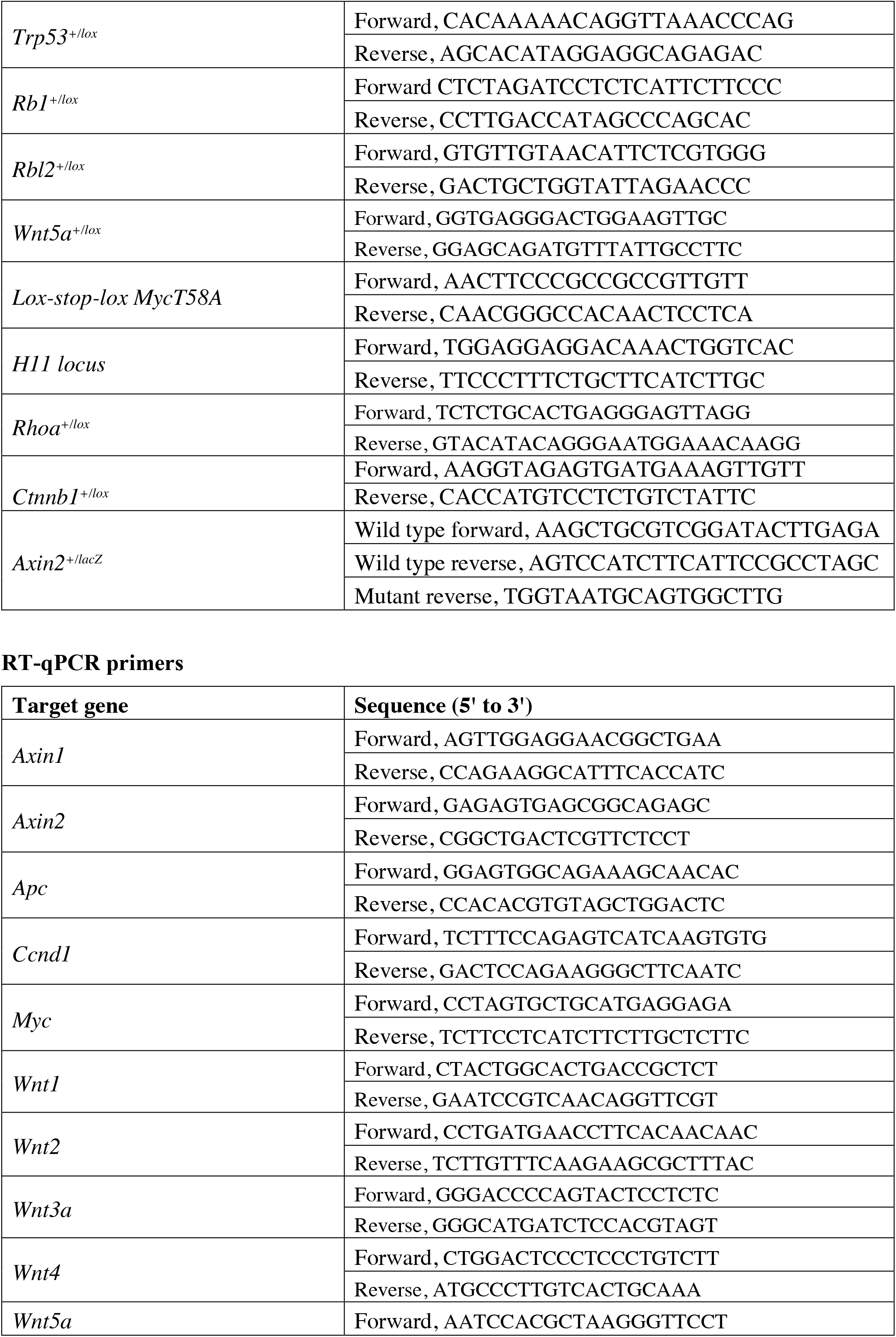
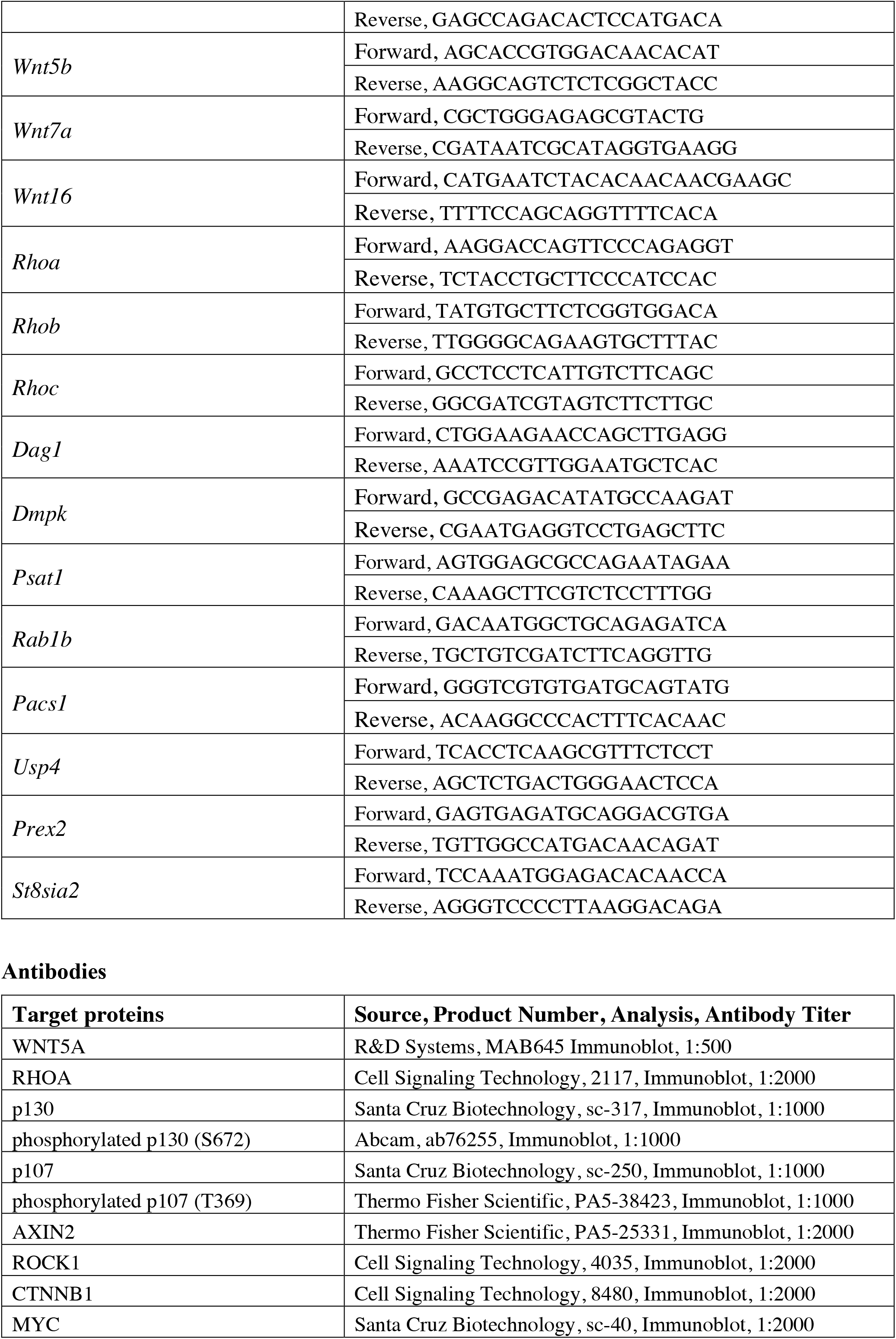
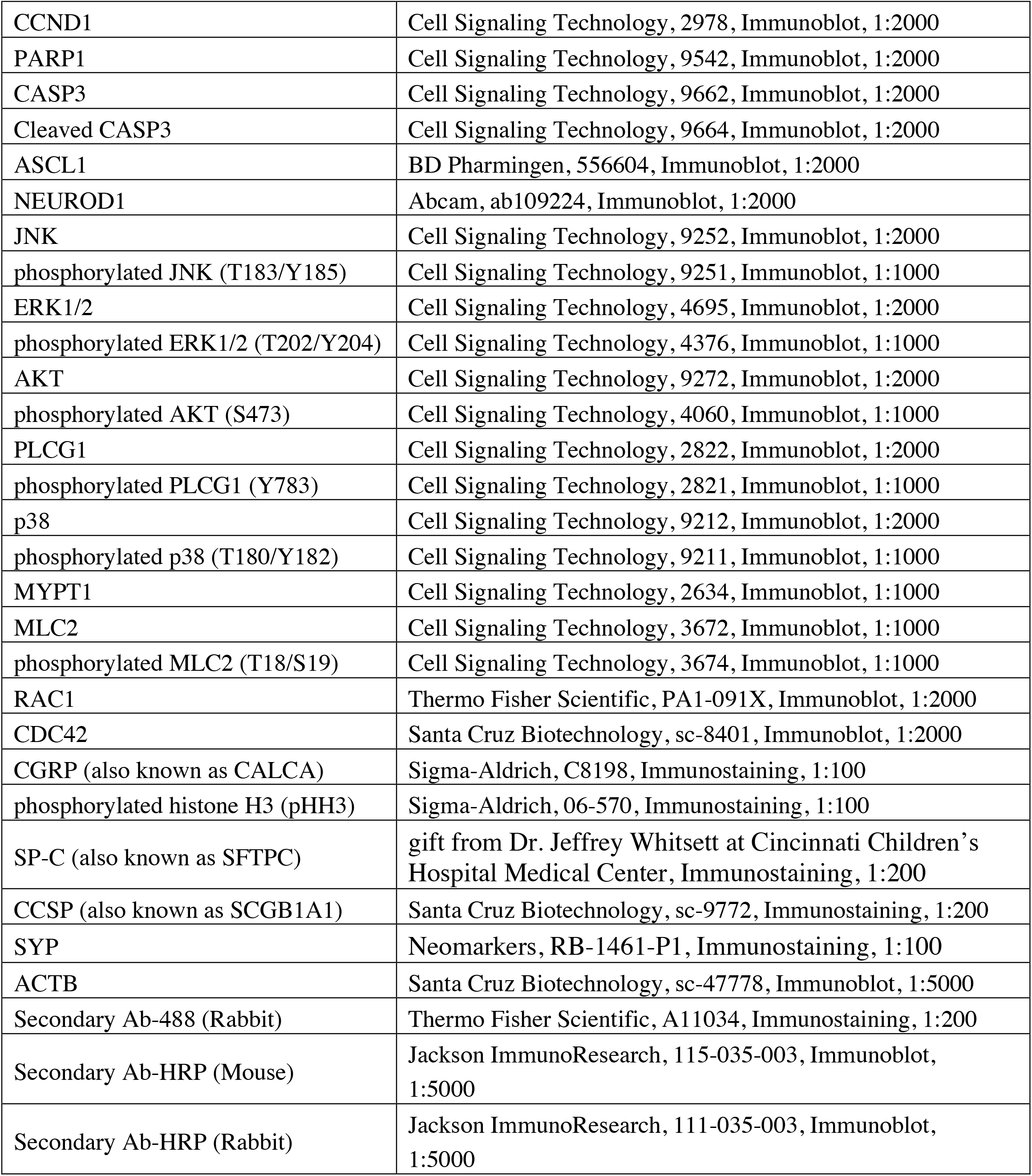
List of plasmids, oligonucleotides, and antibodies used.

